# Autoimmune mechanisms elucidated through muscle acetylcholine receptor structures

**DOI:** 10.1101/2024.12.18.629229

**Authors:** Huanhuan Li, Minh C. Pham, Jinfeng Teng, Kevin C. O’Connor, Colleen M. Noviello, Ryan E. Hibbs

## Abstract

Skeletal muscle contraction is mediated by acetylcholine (ACh) binding to its ionotropic receptors (AChRs) at neuromuscular junctions. In myasthenia gravis (MG), autoantibodies target AChRs, disrupting neurotransmission and causing muscle weakness. Despite available treatments, patient responses vary, suggesting pathogenic heterogeneity. Current information on molecular mechanisms of autoantibodies is limited due to the absence of structures of an intact human muscle AChR. Here, we overcome challenges in receptor purification and present high-resolution cryo-EM structures of the human adult AChR in different functional states. Using a panel of six MG patient-derived monoclonal antibodies, we mapped distinct epitopes involved in diverse pathogenic mechanisms, including receptor blockade, internalization, and complement activation. Electrophysiological and binding assays further defined how these autoantibodies impair AChR function. These findings provide critical insights into MG immunopathology, revealing previously unrecognized antibody epitope diversity and mechanisms of receptor inhibition, offering a foundation for personalized therapies targeting antibody-mediated autoimmune disorders.

**Highlights:**

1. Human adult muscle AChR receptor structures in resting and desensitized states
2. Epitope mapping for mechanistically distinct pathogenic MG autoantibodies
3. Structural and functional elucidation of different inhibition mechanisms in MG

## Introduction

Voluntary movement depends on the contraction of skeletal muscle, a process initiated by the release of acetylcholine (ACh) from motor neurons originating in the brainstem and spinal cord^1^. ACh binds to acetylcholine receptors (AChRs) at neuromuscular junctions (NMJs), triggering ion channel opening, cation influx and depolarization, leading to muscle contraction. Disruptions in this signaling can result in severe muscle disorders^2,3^. A major cause of such disruption is autoantibody-mediated interference, as seen in the autoimmune disease myasthenia gravis (MG)^3,4^. MG autoantibodies targeting the AChR can block neurotransmitter binding, induce receptor internalization, and activate the complement cascade^2,3,5^. This interference leads to muscle weakness and can progress to respiratory failure and paralysis^6,7^.

Current approved MG treatments are focused on increasing ACh availability at the NMJ, broad immunosuppression^2^, and, more recently, complement inhibition, and reduction of circulating immunoglobulin^8-10^. However, the responses to these therapies are variable. For example, 40% of patients respond poorly to complement inhibitors, and in some patients the disease worsens as a result of treatment^9,11^. This variability suggests that the underlying pathogenic mechanisms differ among patients^7,12^. Diagnosis typically involves detecting polyclonal AChR-specific autoantibodies in serum, but this approach overlooks the distinct molecular mechanisms driving MG in individual patients. Moreover, recent findings suggest that single anti-AChR antibodies can act through multiple pathogenic pathways^12^, underscoring the need for a detailed understanding of MG at the receptor-antibody level.

The muscle AChR is the founding member of the pentameric ligand-gated ion channel superfamily^13,14^. It is composed of five homologous subunits: two α1 subunits, and one each of β1, δ and γ (fetal isoform) or ε (adult isoform)^15,16^. These subunits are arranged around a central axis, with an extracellular domain in the synaptic cleft and a transmembrane domain that forms the ion channel pore. The intracellular domain contains binding sites for scaffolding proteins, as well as residues that can tune ion conductance^15,17^. Structural insights into muscle AChRs have been limited, in part due to the complexity of their subunit composition. Most of our knowledge has come from studies of muscle-like ACh receptors from fish electric organs^18-22^, but these models have limited relevance to human diseases^23,24^. Recently, we published the first structures of bovine muscle AChRs, extracting the receptors from bovine muscle^15^. Yet the human adult AChRs remain the gold standard for understanding MG pathological mechanisms. Attempts to express recombinant human muscle ACh receptors have faced longstanding challenges^24-27^. Native AChRs from non-human species exhibit poor cross-reactivity with human diseases^18,23,28^, or are present in quantities unsuitable for characterization of a broad panel of antibody-receptor structures^15,29^.

In this study, we address these challenges by developing a method to express, purify, and determine high-resolution cryo-EM structures of the human adult AChR in distinct functional states. These structures serve as a reference for investigating pathogenic autoantibody mechanisms. We selected a panel of six MG patient-derived autoantibodies, each from a different individual. In addition to representing biological diversity, these autoantibodies were chosen to represent the known mechanisms of MG autoantibody pathology: receptor blockade, internalization, and complement activation. We determined seven high-resolution structures of the Fab fragments individually complexed to the muscle AChR. These provide the first MG autoantibody epitope map for the human muscle AChR, revealing unexpected diversity in antibody binding sites. Complementary electrophysiological and binding assays further elucidate the distinct binding mechanisms underlying pathologic receptor inhibition. Our findings expand the understanding of antibody-mediated autoimmune disorders and MG immunopathology and present novel targets for therapeutic intervention.

### Expression and purification of the adult human muscle AChR

Given the impracticality of obtaining native human muscle AChR in quantities suitable for structural analysis^29^, and the limited relevance of receptors from other species to human disease^18,23,28^, we turned to recombinant expression of the human muscle AChR. This approach has failed to date, due to improper assembly of the receptor when expressed at scale^24,25,27^. To overcome this challenge, we developed a simple and highly reproducible method for recombinant preparation of the human muscle AChR. Specifically, we fused an affinity tag to the δ subunit and paired the α1-δ and β1-ε subunits to minimize incorrect assembly (Figures S1A–D, Methods). Inspired by the natural arrangement of these genes in the human genome^30^—where α1 and δ reside on chromosome 2, and β1 and ε on chromosome 17—we generated a stable cell line that expresses the functional, full-length human muscle AChR. Electrophysiology confirmed that its channel activity is comparable to plasmid-transfected receptors (Figures 1A, 1B). We further optimized expression and purification conditions to produce high-quality receptor preparations (Figures S1F, S1G, S2A, S2B). These samples, combined with optimized cryo-EM processing, enabled high-resolution 3D reconstructions (1.9 to 2.5 Å), with the best map reaching the Nyquist limit (Figures S3, S4). Mass spectrometry and cryo-EM analyses confirmed the presence of only correctly-assembled receptors comprising all four subunit types in the heteropentamer, with no other subunit compositions detected (Figures S2D, S3).

**Figure 1.**
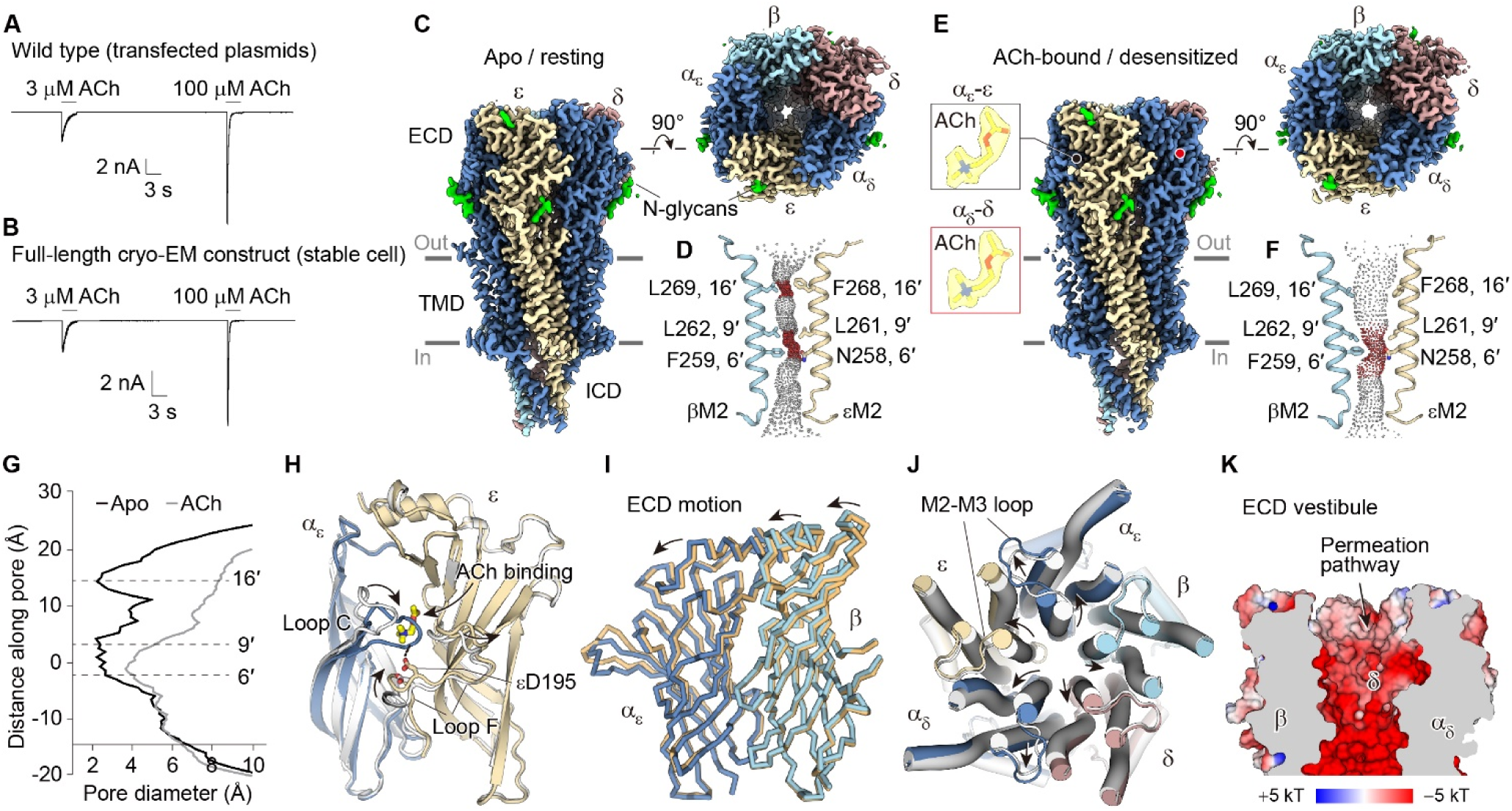
Human muscle AChR structures in resting and desensitized states. (A), (B) Patch-clamp electrophysiology in HEK cells comparing human muscle AChR prepared by transient co-transfections of individual subunits to the stable cell line used for cryo-EM. (C) Cryo-EM density map of receptor with each subunit colored; left panel is side view, upper right panel is top view (90° rotated). N-linked glycans are colored green. (D) Conformation of the transmembrane pore; pore-lining residues of the β and ε M2 transmembrane helices are indicated. (E) Cryo-EM density map of the ACh-bound receptor. Inset is zoom of density (transparent surface) of ACh. (F) Conformation of the transmembrane pore as in D, but for the ACh-bound, desensitized state. (G) Plots of pore diameters comparing apo to ACh-bound states. (H) Conformational differences of the ECD, highlighting movements of the ACh binding site loops, viewed from the receptor periphery. (I) Illustration of global conformational differences between the apo (tan) and ACh-bound (blue) structures, viewed from inside the ECD vestibule. (J) Conformational differences of the TMD helices from top view; apo in gray, ACh-bound in colors. (K) ECD vestibule surface electrostatics.

### Resting and desensitized reference structures

To define reference points for how autoantibodies perturb receptor structure and function, we first sought to determine the structures of apo (unliganded) and ACh-bound human muscle AChRs. We refined the apo and ACh-bound receptor cryo-EM maps both to an overall resolution of 2.1 Å (Figures 1C, 1E, S4). The apo structure serves as a reference for the non-conducting, resting state while the ACh-bound structure represents a non-conducting desensitized state of the human muscle AChR^15,31^. The high-quality density maps enabled *de novo* model building of the receptor as well as ACh molecules, glycans, and lipids (Figure 1E). The structures of human muscle AChR contain five subunits arranged as α_ε_-ε-α_δ_-δ-β around the channel axis (Figures 1C, 1E). Each subunit has an extracellular domain (ECD) composed of mostly β strands. The transmembrane domain (TMD) contains four helices (M1-M4), with M2 lining the channel pore. The intracellular domain (ICD) includes an amphipathic helix (MX) and an MA helix that is continuous with M4. Most features of the human AChR are consistent with that identified in native structures isolated from bovine skeletal muscle^15^, including a long C-terminus stabilized by a disulfide bond in the ε subunit and unique glycosylation sites in the ε and δ subunits, suggesting that mammalian muscle AChRs exhibit high structural similarity.

Comparison of the resting and desensitized state structures reveals asymmetric conformational changes, starting from the ECD upon neurotransmitter binding and transducing into the pore. In both α1 subunits, loop C shifts ∼8 Å toward the neurotransmitter binding site to stabilize ACh (Figure 1H). In parallel, loop F of the complementary subunits (ε and δ) moves towards the closed loop C and stabilizes it by forming a hydrogen bond through a conserved aspartic acid (Figure 1H). Accompanying these local conformational changes, the upper region of each subunit’s ECD undergoes a counterclockwise rotation and bends toward the channel axis, resulting in a ∼3° tilting of the ECD (Figure 1I). Each subunit’s ECD movement alters the components of its coupling region at the ECD-TMD junction to transduce the signal from ACh binding to channel opening^31,32^. Specifically, the M2-M3 loops swing outward, pulling the extracellular ends of M2 away from the channel axis (Figure 1J). Compared to other subunits, β1 undergoes minimal changes in the TMD during the state transition.

The pore conformations between these two states are strikingly different from each other (Figures 1D, 1F). In the resting state, the pore is tightly closed by two extensive hydrophobic gates. The upper gate at 16′ of the M2 helix has a minimal diameter (*d*_min_) of 2.3 Å while the lower gate is located between the L9′ ring and βF6′, resulting in a *d*_min_ at 2.2 Å (Figures 1D, 1G). In the desensitized state, the upper gate widens to 8.5 Å, opening and hydrating the upper half of the channel. By contrast, the lower gate only undergoes a minor increase in diameter (2.2 to 3.7 Å; Figures 1F, 1G) due to the outward movement of M2 helices away from the pore axis.

The L9′ and βF6′ side chains still occlude the permeation pathway, forming the desensitization gate. Although the pore is impermeable to ions in both states, the ECD vestibule is widely open and replete with negative-charges on its inner surface (Figure 1K). We observed several negatively-charged residues along the permeation pathway in the ECD vestibule, including the E20, E49, D85 and D99 residues of the δ subunit. These residues were previously identified as critical for concentrating cations and for cation vs. anion selectivity of the AChR^33,34^. Collectively, these reference structures of the human AChR provide foundational insights into the architecture and conformational changes of the receptor, extending our understanding of how ACh binding stabilizes distinct structural states and offering a detailed framework for investigating autoantibody-mediated perturbations.

### Cryo-EM structures of the AChR bound to pathogenic Fabs

To elucidate the molecular epitopes of autoantibody-mediated pathology, we determined the cryo-EM structures of the muscle AChR bound to individual Fabs at high resolution. Muscle AChR-specific autoantibodies in MG are polyclonal, have varied epitope specificity, and are antigen-driven to undergo affinity maturation and selection^6,7^. We previously generated a library of 11 monoclonal antibodies (mAbs) from the peripheral B cells of ten MG patients^12^. We selected 6 autoantibodies from different patients to sample mechanistic diversity (Figures 2A, 2B). All mAbs are of the immunoglobulin G1 (IgG1) subtypes except for mAb3, which is an IgG3 subtype. Among these autoantibodies, mAb3 exhibits strong blocking of α-bungarotoxin binding to the receptor, which serves as a proxy for neurotransmitter blocking^35^. mAb6 and mAb7 are capable of crosslinking receptors and inducing internalization; and mAb2 and mAb9 activate the classical complement pathway. Some autoantibodies exhibit multiple mechanisms contributing to AChR MG pathology^12^. For example, mAb2 and mAb9 also cause receptor internalization, and mAb1b possesses all three pathological activities. The Fab fragments derived from these mAbs were cloned and purified to form the antigen-antibody complexes in the presence of human muscle AChR (Figures S2G, S2H). Cryo-EM allowed us to obtain seven structures of receptor-Fab complexes that extended to or beyond 2.2 Å resolution (Figures 2C–2I, S4). This high resolution enabled clear mapping of epitope determinants contacted by the variable domains of these Fabs (V_H_ and V_L_ for heavy and light chain variable domains, respectively). Overall, we find that these Fabs exhibit unappreciated heterogeneity in their epitope recognition, their binding stoichiometries, their subunit specificities, and their spatial orientations relative to the receptor (Figures 2C–2I). Different stoichiometries were even found for a single Fab, revealing lower vs. higher affinity sites. All five subunits are involved in complex formation but contribute differentially to autoantibody binding. Most Fabs bind close to the apical surface of the ECD, some sitting on top of the receptor; no Fabs were found near the membrane. In the following sections, we present structural and functional studies on subgroups or individual Fabs organized based on their associated mechanisms.

**Figure 2.**
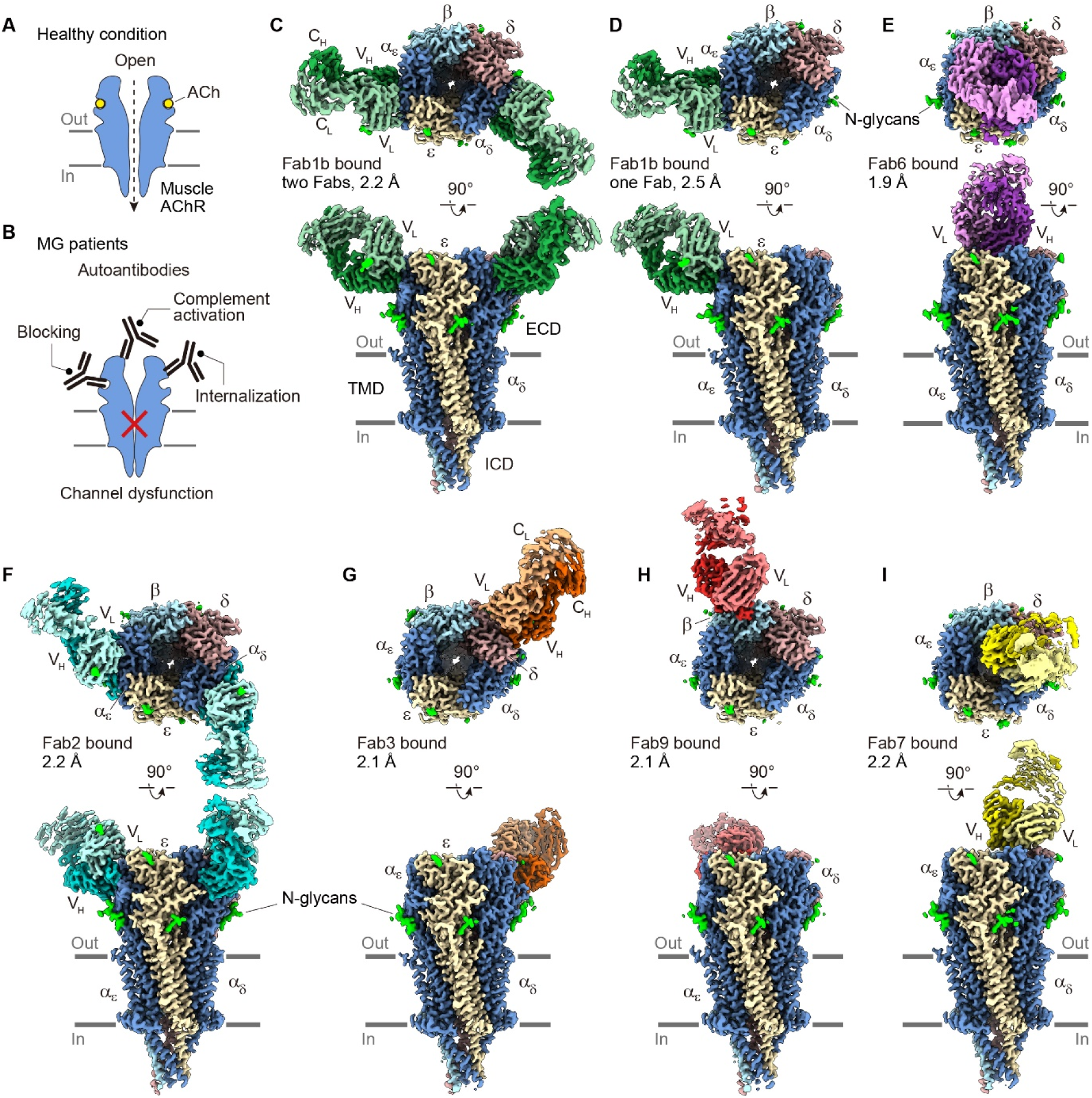
Overview of cryo-EM density maps of myasthenia gravis patient-derived monoclonal antibody fragments in complex with human muscle AChR. (A) , (B) Cartoon illustration of pathological mechanisms of autoantibody interference. (C), (D) Overview maps for the two compositions of Fab1b; one with both α1 subunits bound, the other with only the α_ε_ subunit bound. (E–I) Density maps for the other five antibody-receptor complexes.

### Muscle AChR bound to Fabs from autoantibodies that mediate all three mechanisms

mAb1b was previously found to trigger all three pathogenic mechanisms: blocking, internalization, and complement activation. Our chromatography-based binding assay indicated a 2:1 stoichiometry of Fab1b to the AChR (Figure S2C), suggesting that Fab1b likely binds to the α1 subunits, consistent with cell-based assay results^12^. We determined a 2.2 Å cryo-EM structure of the muscle AChR bound to Fab1b. The structure indeed reveals one Fab1b bound to each of the two α1 subunits, tilted up like ears on a mouse, at an angle of ∼60° relative to the channel axis (Figures 3A, S8B). Extensive electrostatic interactions involving residues in V_H_, V_L_, and the receptor are observed. R139 of the V_H_ complementarity-determining region (CDR) 3 is positioned to form a salt bridge and an additional hydrogen (H) bond with the E23 side chain on the α1 subunit (Figure 3B). The V_H_ CDR3 Q132 inserts deeply, contacting the backbones of αP21 and αE23 to form H-bonds. Several residues on V_H_ CDR1 and CDR2 also form multiple H-bonds with the α1 subunits to stabilize Fab1b binding (Figure 3C). The interaction patterns on α_ε_ and α_δ_ subunits are similar to each other, however, the epitopes contacted by the V_L_ of Fab1b are different at the two complementary subunit (ε/δ) interfaces (Figure 3D). In the α_ε_-ε interface, K52 from V_L_ CDR3 is tightly bound via electrostatic interactions with two bulky polar residues, εK69 and εE76, from the top of the ε ECD. By contrast, the corresponding region in the δ subunit harbors a large glycan at δN76, which restricts access of the Fab1b light chain. Structural comparison reveals that the δ ECD is pushed away from the α_δ_ subunit to accommodate Fab1b binding because of the limited space (Figure 3D, dashed arrow). In addition, in the δ subunit, these two charged interacting residues are alanine and serine respectively. Thus, the two binding sites for Fab1b are not equivalent, highlighting differences in epitope specificity for the same Fab.

**Figure 3.**
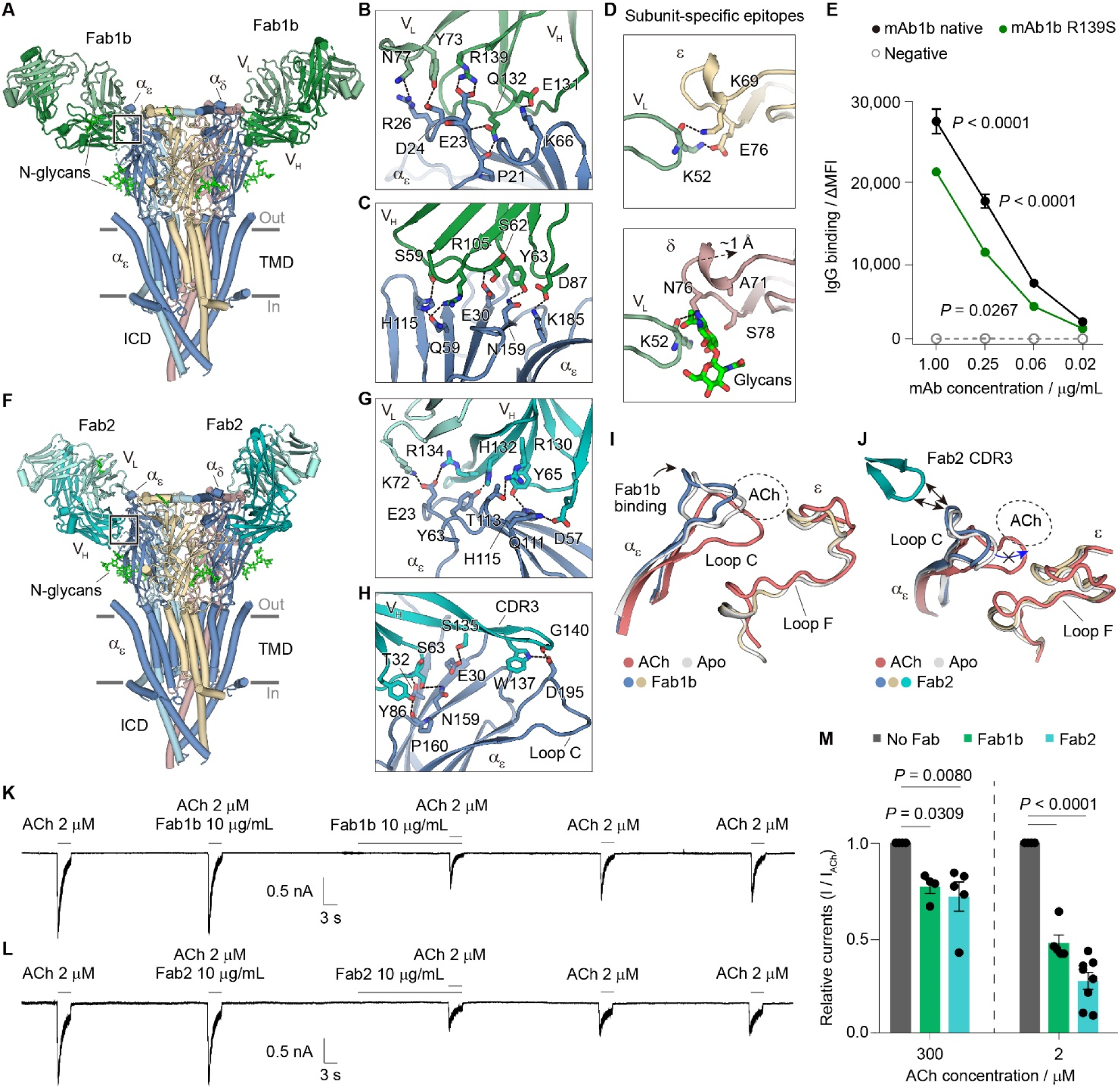
Structural and functional interrogation of the Fab-AChR complexes targeting the α1 subunit. (A) Overview cartoon of Fab1b-receptor complex structure with insets in following panels indicated. (B), (C) Insets focused on interacting residues between Fab1b (V_H_, dark green; V_L_, light green) and receptor (blue). Dashed lines indicate electrostatic interactions. (D) Fab1b light chain reveals different epitopes between ε and δ subunits of the AChR. (E) Titration plot binding curves of the native and mutant mAb1b tested for binding over a range of serially diluted concentrations. Negative control, mAb58 (aquaporin-4 specific). Y-axis is difference in normalized mean fluorescence intensity (MFI) of secondary antibody vs. MFI with primary mAb + secondary antibody. Data are expressed as mean ± s.e.m. of experimental triplicate values. (F) Overview of the Fab2-receptor complex with insets in panels G, H indicated. (G), (H) as for B, C but Fab2 is in teal. (I), (J) Fab1b and Fab 2 binding lead to a more opened loop C in the ACh binding site. (K), (L) Representative traces of inhibition of ACh-induced currents by Fab1b or Fab2 by whole-cell patch-clamp electrophysiology. (M) Quantification of inhibition of 2 M or 300 M ACh-induced currents by Fab1b or Fab2; related to Figure S5. Data points indicate the biological replicates; mean ± s.e.m.

Interestingly, we observed a small fraction of particles (18%) with only one Fab1b bound per receptor, despite the major 2:1 stoichiometry in the dataset (Figures 2C, 2D). In this minor population, Fab1b only binds to the α_ε_ and ε subunits through interactions similar to those in the two Fab-bound structure, illustrating how subtle epitope changes influence antibody affinity.

To understand how individual residues of mAb1b contribute to binding the receptor, we mutated residues that made electrostatic and hydrogen-bond contacts (shaded residues, Figure S6A). We also focused on residues that had evolved as a result of somatic hypermutation (SHM), a process that is unique to the immunoglobulin genes of B cells and is known to select for increased affinity of antibodies to their targets^6,36^.

Such SHM residues were mutated “back” to the original germline-encoded amino acid as predicted by alignment to the IMGT database^37^. Non-SHM residues were simply mutated to alanine. Most individual mutations did not significantly decrease binding (data not shown). However, R139 of the heavy chain CDR3, which makes a salt bridge with E23 of the α_ε_ subunit, is clearly important for antibody binding (Figure 3E).

The mAb2 antibody was cloned from a different patient and did not block α-bungarotoxin binding, but had dual functionality in complement activation and internalization. As for Fab1b, it is also predicted to have a 2:1 stoichiometry (Figure S2C). We determined the structure of the muscle AChR bound to Fab2 at a 2.2 Å resolution (Figures 2F, 3F, S4). Interestingly, we only identified particles bound with two Fabs in the dataset; no single Fab-bound particles were found, likely because Fab2 exclusively binds to the α1 subunits (Figures 3F–H), consistent with cell-based assay results^12^. Thus, the two binding sites for Fab2 on the muscle receptor are equivalent. Fab2 has a similar spatial binding angle (∼65°, as measured from the central axis of the receptor, Figure S8C) to Fab1b but shifts ∼33° clockwise around channel axis to move away from the two complementary subunits (Figure S7A). Although their binding positions partially overlap, most epitope determinants for Fab2 and Fab1b are distinct. We only observed one H-bond interaction from the Fab2 V_L_; the extensive electrostatic interactions with the receptor involved the Fab2 V_H_ (Figures 3G, 3H, S6). Interestingly, the Fab2 V_H_ CDR3 folds into a long pair of β-strands, extending from one side of the α1 subunit to reach its loop C on the other side (Figure 3H). In particular, W137 and G140 in the tip of the CDR3 β-strands form H-bond interactions with D195 on the tip of α1 loop C.

Comparisons of Fab1b- and Fab2-bound structures with the two reference structures reveal that both Fabs stabilize a more open conformation of loop C, which normally should close upon ACh binding. Fab1b closely contacts the receptor, forcing a slight outward rotation of the α1 ECD as a rigid body, which in turn causes loop C opening (Figure 3I). Interestingly, Fab2 binding also leads to the α1 loop C opening slightly compared to the apo structures, through direct interactions with its CDR3 (Figure 3J). In both cases, locking loop C open should logically destabilize ACh binding, which requires loop C to pack around it for high affinity, and to trigger channel opening. To test this hypothesis, we performed whole-cell patch-clamp recordings on our stable cell line. Both Fabs decreased ACh-dependent channel responses at both high and low ACh concentrations (Figures 3K, 3L, S5). Furthermore, both Fabs washed off slowly, consistent with their extensive interactions with the receptor.

Together these findings reveal that both Fab1b and Fab2 interfere with muscle receptor activity by destabilizing ACh binding, although they have different subunit epitope determinants, arise from different immunoglobulin genes, and are from different patients^12^.

### Mechanism of direct competition with ACh by Fab3

Of the patient-derived autoantibodies screened in our initial study, mAb3 is the most effective at competing with α-bungarotoxin binding to the receptor^12^. We found that Fab3 could not bind to the α-ε pentameric receptor (Figure S2D), suggesting that it attacks the α_δ_-δ interface. To understand its inhibition mechanism at the atomic level, we determined the structure of muscle AChR bound to Fab3 at 2.1 Å resolution (Figures 2G, 4A, S4). The structure reveals that Fab3 binds with a spatial angle at ∼62° to the receptor in a ratio of 1:1, and the majority of its contacts are with the δ subunit, with minor contacts on the β1 and α_δ_ subunits (Figures 4A, S8D).

**Figure 4.**
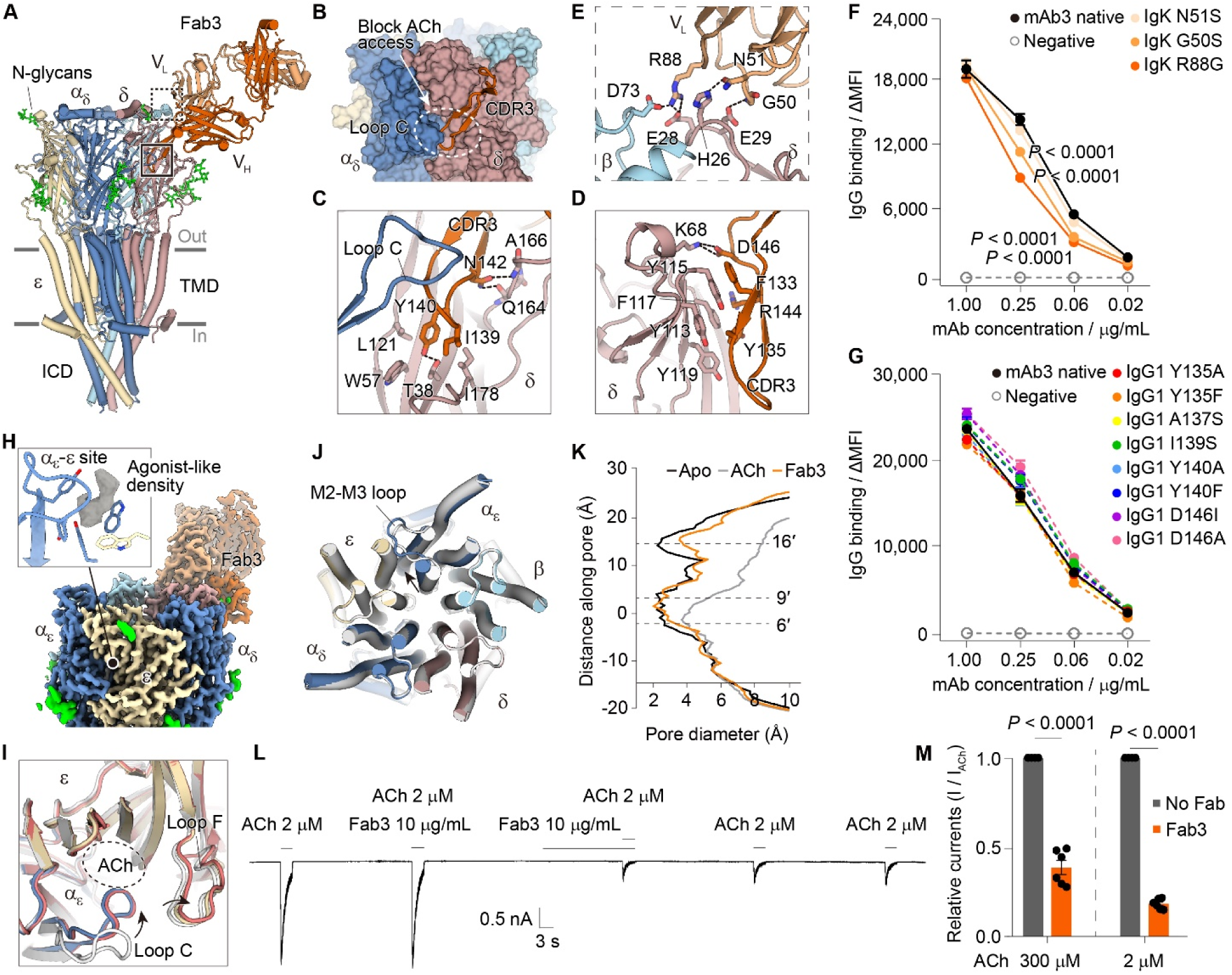
Competitive AChR antagonism by Fab3. (A) Overview cartoon of Fab3-bound AChR complex with insets indicated in boxes. (B) Illustration of direct competition by Fab3 V_H_ CDR3 (ribbon) in the ACh binding site of AChR (surface). (C), (D) Interacting residues involved in B are illustrated from two vantage points to highlight all contributions from CDR3 to binding the δ subunit. (E) Light chain residues that are critical for interaction are shown; dashed lines represent electrostatic interactions. (F), (G) Titration plot binding curves of the native mAb3 and its mutants tested for binding over a range of serially diluted concentrations, as for Figure 3E. F, light chain mutants; G, heavy chain mutants. Data are mean ± s.e.m. (H) Density for a potential agonist in the ACh site of α_ε_-ε interface. (I) Comparison of loop C closure between the α_ε_-ε interfaces; apo (gray), ACh-bound (red), Fab-3 bound (blue and tan). Note this interface is not the one occupied by Fab3. (J) Comparisons of the conformations of TMD helices between apo and Fab3-bound state. (K) Comparisons of pore diameters. (L) , (M) Functional inhibition and quantification of inhibition of ACh-induced currents by Fab3 measured by whole-cell patch-clamp electrophysiology. Data points indicate biological replicates; mean ± s.e.m. Related to Figure S5.

The extraordinary length of the V_H_ CDR3 loop, 29 amino acids, allows it to reach from the δ subunit all the way into the neurotransmitter site at the interface of α_δ_ and δ subunits (Figure 4B, Video S1). In contrast to Fab1b and Fab2, the Fab3 V_H_ binds to the receptor mainly through hydrophobic interactions (Figures 4C, 4D). Two residues, I139 and Y140, along with the tip of V_H_ CDR3 insert into the proximity of the ACh binding pocket (Figure 4C), where they are embraced by several hydrophobic residues from the ACh binding site. This insertion limits ACh access to its pocket and sterically hinders the movement of loop C to inhibit ACh binding (Figure 4B). Several π-π and cation-π interactions were observed between the V_H_ CDR3 and a hydrophobic residue cluster on the δ ECD (Figure 4D). This pattern explains the δ-subunit specificity of Fab3, which cannot bind to α_ε_-ε interface where the hydrophobic cluster has been replaced by small polar resides in the corresponding region of ε ECD (Figure S7B). Notably, we also observed an electrostatic interaction network, mediated by the R88 on V_L_ and anionic residues from both δ and β subunits to stabilize the Fab3 light chain binding (Figure 4E).

SHM analysis reveals that both Fab3 V_H_ and V_L_ contain replacement residues involved in receptor binding (Figure S6). Of particular interest, we found that a glycine mutates to an arginine (R88) to stabilize the V_L_ binding; this large side chain change motivated us to evaluate its importance for mAb3 maturation. We found that indeed, mutation of R88 to the germline glycine decreased binding significantly (Figure 4F). So did mutation of the nearby G50 to serine, which we hypothesize decreases flexibility of the backbone and prevents backbone interactions with E29 of the receptor δ subunit (Figures 4E, F). In contrast, we did not see any effect from individual mutations of V_H_ CDR3 hydrophobic resides on binding (Figure 4G), suggesting that these smaller interactions contribute as a dispersed hydrophobic network to bind the receptor.

Interestingly, we noticed an agonist-like density in the ACh site at the α_ε_-ε interface, which resembles an ACh molecule (Figure 4H). We did not model this density as no agonist was included during sample preparation, and thus we are not confident in its identity. Binding of this agonist-like molecule in the α_ε_-ε interface leads to α_ε_ loop C closure, similar to that observed in our ACh-bound desensitized reference structure (Figure 4I). However, only the M2-M3 loop and the extracellular end of M2 extracellular end of the α_ε_ subunit are pulled outward from the channel axis, while the other subunits maintain conformations identical to the apo-resting reference structure (Figure 4J). These changes result in an impermeable channel with a slightly-enlarged upper gate (*d*_min_ at 3.6 Å) at M2 16′, which resembles a resting-like pore conformation (Figure 4K). These data suggest that Fab3 acts like a wedge in the α_δ_-δ interface, blocking ACh binding and preventing channel activation even while agonist-like molecules may bind freely to the second ACh site at the α_ε_-ε interface. We evaluated the inhibition by Fab3 in patch-clamp electrophysiology (Figures 4L, 4M, S5). Our results show that Fab3 efficiently inhibits the channel response at both high and low concentrations of ACh, and washes off very slowly, consistent with its extensive hydrophobic and polar interactions with the receptor. These findings indicate that Fab3, acting as a strong antagonist, inhibits the receptor by stabilizing a resting-like state.

### Fabs sitting on the muscle AChR apex

Among our patient mAbs, mAb6 and mAb7 predominantly induce receptor crosslinking and internalization^12^. We previously could not determine their subunit specificities, suggesting a unique binding action for these autoantibodies compared to those dominated by the α1 or δ subunits. To understand their subunit specificity and pathology, we determined the structures of muscle AChR bound to Fab6 and Fab7 at 1.9 Å and 2.2 Å resolution, respectively (Figures 2E, 2I, 5A, 5F, S4). To our surprise, in contrast to other Fabs, both Fab6 and Fab7 bind to the very “top” of ECD of the receptor and attack multiple subunits.

**Figure 5.**
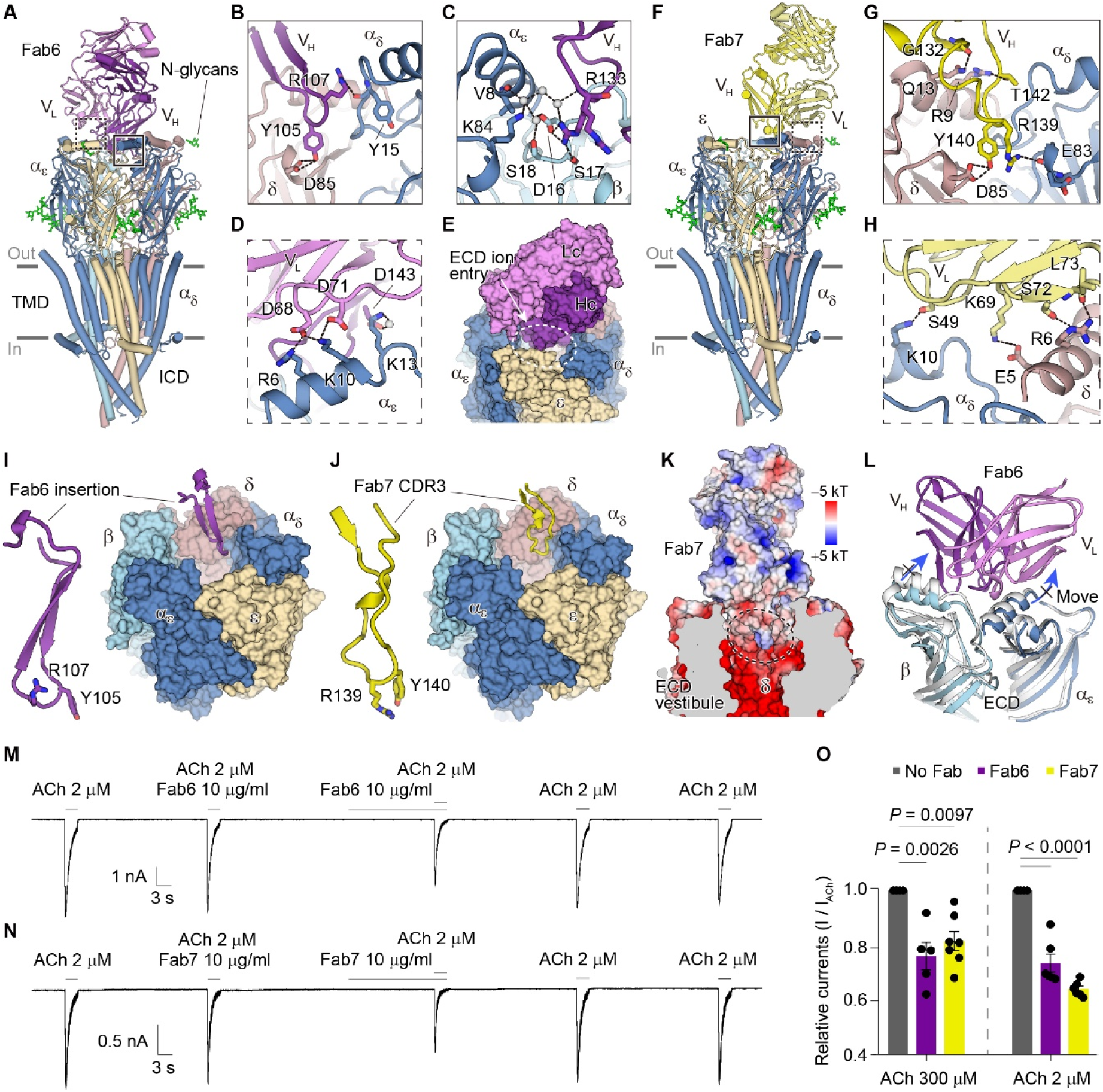
Structural and functional interrogation of the internalization-competent Fab6 and Fab7 antibodies bound to AChR illustrate occlusion of the extracellular vestibule. (A) Overview cartoon of Fab6 binding to muscle AChR. (B) Residues of Fab6 V_H_ contact both the δ and α_δ_ subunits. Dashed lines indicate electrostatic interactions. (C) A coordinated network of water molecules (small grey spheres) contributes to binding of Fab6. (D) Fab6 light chain residues contribute to α_ε_ subunit binding. (E) Surface model of Fab6-AChR seen from the perspective of the synaptic junction. Dashed oval indicates proposed entry of ions into the vestibule. (F) Overview cartoon of Fab7-AChR complex. (G) , (H) Residues from the V_H_ and V_L_ of Fab7 make electrostatic interactions with the α_δ_ and δ subunits. (I) Fab6 and (J) Fab7 loops insert into the ECD vestibule and occupy similar sites on the δ subunit, engaging D85 and increasing the electropositivity of the vestibule (K). (L) Fab6 binding prevents conformational contraction of the ECD seen in the ACh-bound desensitized state. (M–O) Fab6 and Fab7 functionally inhibit maximal ACh current responses in whole-cell patch-clamp electrophysiology. Data points in O indicate biological replicates; mean ± s.e.m. Related to Figure S5.

Fab6 binds nearly on the long axis of the ECD, with a ∼34° tilt towards the ε subunit, and contacts all four subunits except ε (Figures 5A, 5E, S8F). However, only a few electrostatic interactions between both V_H_ and V_L_ and the receptor were observed, consistent with its weaker binding strength compared to other mAbs^12^. The most striking feature is that Fab6 binding to the ECD apex leaves only two small entries for cation influx, restricting access and impeding the ability of the ECD vestibule to concentrate cations (Figure 5E). Two bulky residues, Y105 and R107 in V_H_, insert deeply into the ECD vestibule (Figure 5B). Y105 forms a hydrogen bond with the negatively-charged D85 on the δ ECD inner surface, while R107 contacts the residue backbone on the α_δ_ subunit. Because of the high structural resolution, we were able to observe alternative side-chain rotamers and several well-ordered water molecules that mediate the interaction networks (Figure 5C). For example, one rotamer of R133 on V_H_ CDR3 interacts with a negatively-charged residue, D16 on the β1 ECD, and its adjacent waters to stabilize the Fab6 V_H_ binding. The α_ε_ ECD is also involved, where R6 and K10 on the α_ε_ ECD contact anionic residues to stabilize the V_L_ binding (Figure 5D). All of these interactions contribute to the novel function of sterically “plugging” the entry of ions into the extracellular channel vestibule. Similarly, Fab7 predominantly stands on the δ subunit, with a spatial angle of ∼10°, and also extends interactions to a few residues on the α_δ_ and β1 subunits (Figures 5F, S8G). In an example of convergent evolution across two patients, Fab7 uses two bulky residues-R139 and Y140, within V_H_ CDR3, to insert deeply into the ECD vestibule (Figures 5I, 5J). There, Y140 of Fab7 interacts with the negatively-charged residue D85 on the δ ECD inner surface through hydrogen bonds, while R139 contacts the backbone in the α_δ_ subunit (Figure 5G). The V_L_ of Fab 7 also contributes by forming several electrostatic interactions with the polar residues on the δ and α_δ_ subunits (Figure 5H).

Although Fab6 and Fab7 arose from different gene ancestors (Fab6: IGHV 3-23/D4-23*01/J4*02; Fab7: IGHV 3-74/D3-16*01/J5*02) and originate from two different patients, they appear to have a convergent mechanism to interfere with muscle receptor function. Both the Fab6- and Fab7-bound structures harbor an extended loop in their heavy chains, contacting the same negatively-charged residue and introducing positive charges into the vestibule surface (Figure 5K), which is detrimental for concentrating cations in the ECD vestibule^31,33,34,38^. These interacting residues in the extended loop arose from an insertion of N nucleotides at the VD junction in mAb6, and an insertion of N nucleotides between the DJ junction in mAb7. Furthermore, Fab binding on top of the ECD pushes the contacted subunits outward from the channel axis. Structural comparison with the desensitized reference state reveals that Fab6 binding would hinder the contractile movement of ECD induced by ACh binding (Figure 5L). These analyses suggest that Fab binding here alters the ECD vestibule net charge and prevents conformational changes in the receptor, both of which can impair channel activity (Figure 5K). Consistently, electrophysiology assays reveal that both Fabs significantly reduce ACh-dependent channel responses at both high and low ACh concentrations (Figures 5M–5O, S5). These findings reveal that in addition to receptor internalization, mAb6 and mAb7 interfere with muscle receptor channel activity by neutralizing ECD vestibule electrostatics and by blocking receptor state transitions.

### Fab9 binds to β subunit MIR-like region

In its monoclonal IgG form, mAb9 can activate complement and cause internalization through receptor crosslinking^12^. We show that Fab9 has a 1:1 binding ratio to the muscle receptor and shifts the β1 homopentamer elution volume in size-exclusion chromatography, in line with previous cell-based assay results showing that it mainly binds to the β1 subunit^12^ (Figures S2C, S2E). We determined the receptor structure bound to Fab9 at 2.1 Å resolution (Figures 2H, 6A, S4H). The complex structure shows that Fab9 indeed exclusively contacts the β1 ECD outside surface, with a spatial angle at ∼70° (Figure S8H). Interestingly, we found that the V_H_ CDR3 reaches into a region on the β1 subunit homologous to the “main immunogenic region” (MIR) of α1 (Figure 6C). There, β1 D68 mediates an electrostatic network with R139 on V_H_ and Y68 on V_L_ while β1 H72 interacts with V_H_ D137 through a H-bond. In addition, V_H_ CDR1 and CDR2 also participate in formation of the extensive polar interactions with β1 ECD (Figure 6B). For example, V_H_ R89 forms a salt bridge and H-bond with β1 D27; both V_H_ Y66 and E133 are arranged to form H-bonds with β1 R23, all of which contribute to stabilize the strong binding. Despite mutating E133D, D137A, and Y68A, none of these individual mutations contributed significantly to binding, suggesting that Fab9 interaction depends on a distributed network of interactions rather than a key salt bridge or residue (Figures 6B, 6C, S7C).

**Figure 6.**
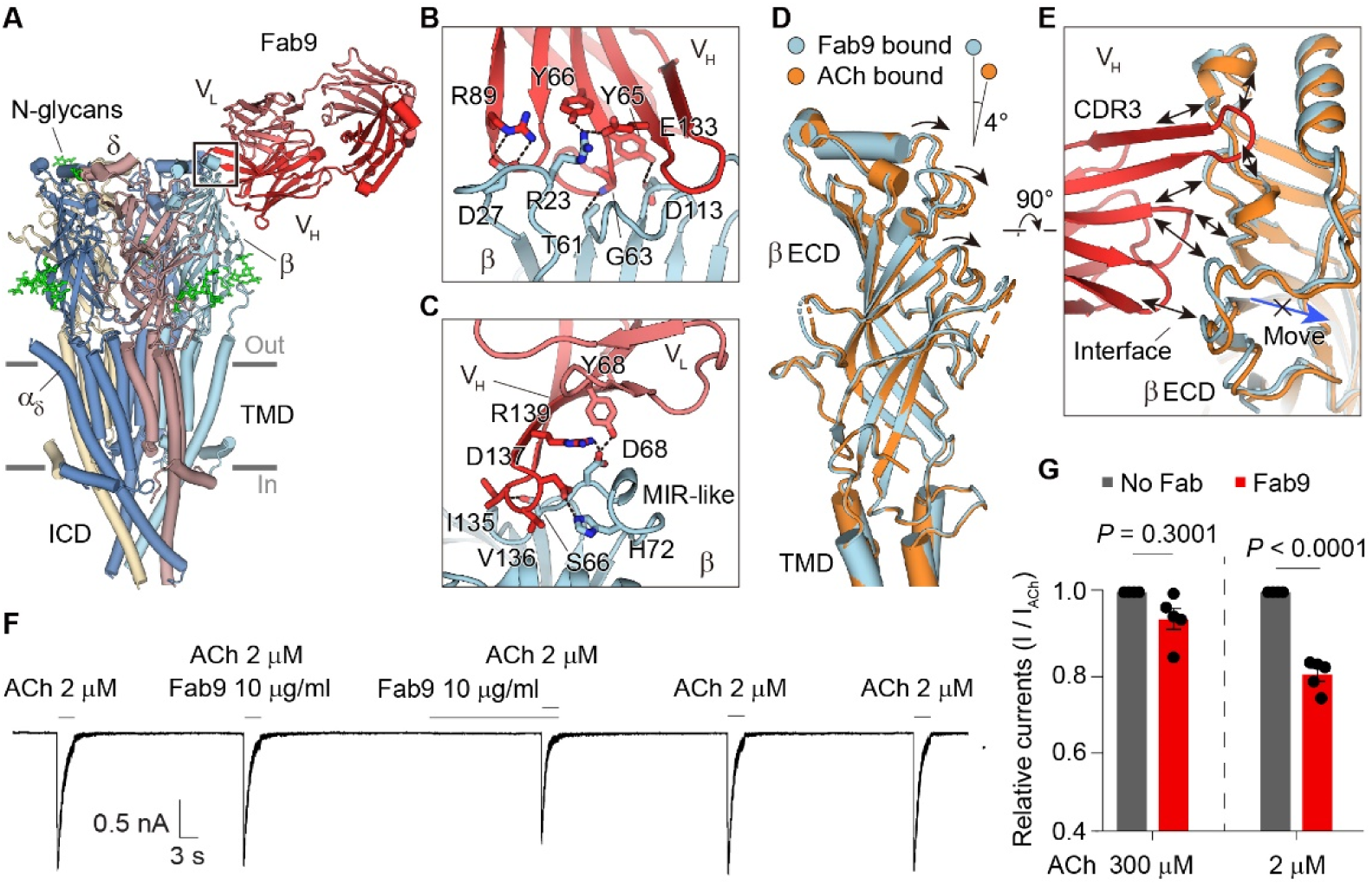
Structure and electrophysiological analysis of Fab9-AChR interactions. (A) Overview of Fab9-AChR structure highlighting angle of binding of Fab9 to the β1 subunit. (B), (C) Residue interactions between the heavy and light chains of Fab9 on β1, including contacts with a region analogous to the MIR. (D), (E) Conformational differences in the ECD between the Fab9-bound (blue) and ACh-bound structures (orange) reveal that Fab9 binding prevents conformational transition. (F) Functional inhibition by Fab9 on ACh response of AChR in whole-cell patch clamp electrophysiology. (G) Quantification of inhibition of ACh response by Fab9. Related to Figure S5. Data points indicate the biological replicates; mean ± s.e.m.

The receptor ECD undergoes contraction and tilting upon binding of ACh to transduce signals to the pore (Figure 1I). Comparison of the resting and desensitized reference structures reveals that the β1 ECD must tilt 4° to respond to ACh (Figure 6D). However, this local ECD movement is prevented by the presence of Fab9 (Figure 6E). This subtle inhibition of ECD movement by Fab9 could lead to a reduction of receptor function, and thus we evaluated the effect of Fab9 by patch-clamp recordings. The results show that the Fab9 inhibition is negligible at a high concentration of ACh, but it indeed decreases ACh-dependent channel responses to a low ACh concentration (Figures 6F, 6G, S5). These data suggest that while Fab9’s major pathological functions are likely complement activation and receptor internalization, it can also interfere with channel activity by preventing conformational transitions between different states. The physiological relevance of a lower concentration of ACh may be significant during neurotransmission as it is rapidly degraded by acetylcholinesterase in the NMJ. Thus, residual channel openings of the receptor would be prevented in the context of Fab9 binding.

### Human muscle AChR epitope mapping

AChR autoantibodies are polyclonal and can bind to multiple subunits^6,7,12,39^. Looking at all six Fab structures, we observed high heterogeneity in spatial binding position, subunit specificity and epitopes. To understand overall binding positions, we merged all structures into one to generate a comprehensive binding map of these pathogenic autoantibodies on the muscle receptor surface (Figure 7). Notably, Fab6 and Fab7 overlap in part in their binding positions, and the same is true for Fab1b and Fab2, although they have different epitopes (Figure 7A). We found that the α1 subunit harbors more epitope-determining residues than β1, δ and ε, consistent with previous findings^40,41^ (Figures 7B, 7C). Interestingly, δ and β1 subunits contain abundant epitope determinants, approaching the number from the α1 subunit. They do have less frequency, likely due to the two copies of α1 subunit per receptor versus one each for δ and β1. The ε subunit only contributes two residues to epitopes, resulting from the extensive reach across the α_ε_-ε interface by Fab1b (Figures 7B, 7C). We found that in each subunit, most epitope determinants are located on the outside surface of the ECD, focusing on a position composed of β-strands and nearby loops. Because of the deep insertions of Fab6 and Fab7, several determinants are also present in the inner ECD vestibule (Figure S7D). Although most Fabs have interactions with either or both α1 subunits, the α1 MIR region played no role in binding of any of these MG-derived Fabs (Figure 7B). By contrast, we observed several epitope determinants on the β1 MIR-like region, stemming from Fab9 V_H_ CDR3 interactions (Figure 6C).

**Figure 7.**
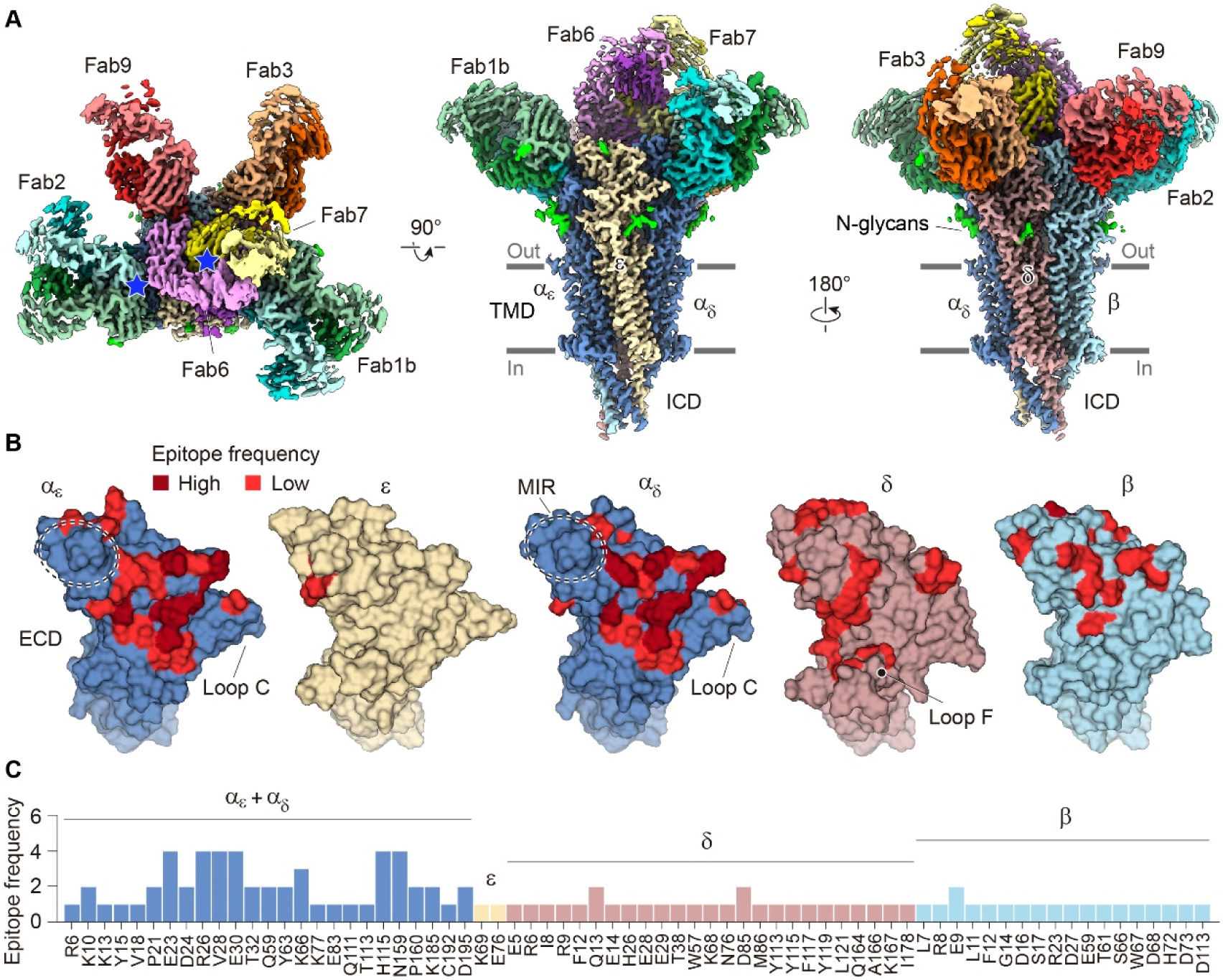
Epitope mapping of MG autoantibodies on the high-resolution map of the human muscle AChR. (A) Views from the synaptic cleft and two side views illustrate overlapping (blue stars) and distinct binding epitopes across patients. (B) Surface representations of the extracellular domains of individual subunits are color-coded by antibody interactions mapped by cryo-EM. Notably, the MIR of the α1 subunits is not a part of the epitopes determined in these structures. Only epitopes in the ECD outside surface are shown here; epitopes in the vestibule inner surface are shown in Figure S7. (C) Frequency of epitopes on receptor subunits contacted across the panel of six patient-derived structures.

## Discussion

### An extended concept of immunopathology

AChR autoantibodies in MG are thought to cause dysfunction in neuromuscular junction signaling through up to three distinct mechanisms, resulting in the characteristic muscle weakness in patients^2,42^. One proposed mechanism, competition with the neurotransmitter acetylcholine, has traditionally been based on indirect measurements. Specifically, MG autoantibodies—polyclonal or monoclonal—are tested for their ability to compete with the binding of radiolabeled α-bungarotoxin^43,44^. This peptide from snake venom binds the two ACh sites on the receptor with sub-nanomolar affinity, inducing paralysis^18,45,46^. This assay is highly specific due to the exquisite selectivity of α-bungarotoxin for the AChR. However, a key limitation of this diagnostic assay is that MG autoantibodies may perturb ACh binding and channel activation without interfering with α-bungarotoxin binding^12^. As a result, false-negative outcomes in this assay have contributed to the perception that most MG autoantibodies do not compete with ACh binding and thus do not directly affect channel function^47^. Structural and electrophysiological studies circumvent this limitation by directly visualizing the inhibition of ACh binding and measuring the receptor’s functional response to ACh in the presence of pathogenic MG antibodies or their fragments.

By combining structural and electrophysiological approaches, we propose a revision to the prevailing view that direct inhibition of the receptor is rare compared to mechanisms like crosslinking and complement activation in MG patients^48,49^. Our findings show that all six MG-derived Fabs inhibit receptor function to varying degrees, suggesting that the role of direct inhibition may be underestimated^2,50^. This conclusion aligns with recent electrophysiological studies showing that sera from 6 of 11 patients caused an immediate reduction in ACh-induced currents in cells expressing the human muscle AChR^50^. The precise mechanism by which autoantibodies directly influence receptor function depends upon their specific binding sites. Binding to the receptor is intrinsic to the pathology, and in theory, the entire extracellular domain (ECD) of the muscle AChR is a potential target for autoantibody attack. The ACh binding site accounts for only a small portion of this surface, meaning that previous assumptions focused solely on blocking ACh binding may overlook other possible mechanisms of interference.

To refine our understanding of MG pathology, we propose expanding the concept of "blocking ACh binding" to "interfering with channel activity". Using structural analysis and electrophysiology, we identified at least four distinct ways by which antibodies can disrupt normal channel function: 1) Direct competition with ACh binding. For example, Fab3 inserts its CDR3 loop in a way that physically blocks ACh access. 2) Destabilizing ACh binding. Fab1b and Fab2 prevent loop C closure, destabilizing ACh’s interaction with the receptor. 3) Neutralizing the receptor’s cation-concentrating potential. Fab6 and Fab7 alter surface electrostatics by forming salt bridges in the receptor vestibule, which influences cation influx. 4) Preventing receptor conformational transitions. Both Fab6 and Fab9 bind in a way that allosterically limits the receptor’s conformational changes required for ACh binding and channel activation. These mechanisms are closely linked to the spatial binding positions of the autoantibodies, with some likely exerting effects through multiple mechanisms. Autoantibodies causing GABA_A_ and NMDA receptor encephalitis have also recently been shown to cause competitive and allosteric antagonism of their respective ion channel targets^51,52^, demonstrating broad relevance of channel inhibition in autoantibody-mediated diseases.

### Structural implications for different pathologies

Structural analyses can aid in predicting the pathogenic mechanisms of MG autoantibodies. For example, in the classical complement pathway, six antibodies need to interact with each other by assembling a hexagonal Fc platform for C1q recruitment and signaling activation (Figure S8A)^53,54^. There, Fabs are arranged to form an ordered array with optimal spatial angles to bind clusters of surface antigens. In the IgG1-C1 initiation complex^53,54^, structural constraints between Fabs and Fc only allow monovalent binding of each antibody (Figure S8A). Superpositions of an intact IgG1 antibody (PDB 1IGY)^55^ onto our Fab complex structures suggest that most of those tested in this study bind in a spatial angle ranging from ∼60° to ∼70° from the central axis of the receptor. This angle projects the IgG Fc toward a position where it could feasibly connect with other Fc of the hexamer as well as present Fc residues required to bind C1q (Figure S8). This finding suggests that these angles are suitable for hexameric Fc formation of IgG1 and C1q binding. Consistent with this hypothesis, we previously found that the autoantibodies in our study that show an angle within this proposed range (mAb1b, 2 and 9) are able to efficiently activate complement^12^. By contrast, mAb6 and mAb7 have a much smaller spatial angle (below ∼30°), which may introduce too large of structural constraints to form an IgG1 hexamer even though the antibody hinge flexibility is considered, and therefore would inefficiently recruit C1q^12^. Interestingly, although mAb3 has a suitable angle, the orientations of its light and heavy chains undergo a rotation compared to other Fabs, for example Fab9, which may impair the optimal antibody platform assembly (Figure S8E). However, as mAb3 is natively IgG3, the extended hinge region associated with this immunoglobulin subtype likely compensates for complement activity that is observed with this autoantibody. Thus, both the spatial binding angle and precise orientation of an MG antibody are important for its ability to activate complement. The binding orientation and proximity have also been shown to be essential for IgG1 antibodies in other autoimmune diseases, such as in neuromyelitis optica, where aquaporin-4 autoantibodies have to be well-organized to enhance optimal Fc-Fc interactions for efficient complement activation^56-58^.

Structural modeling of the MG autoantibodies here suggests that geometric and steric limitations do not allow two Fabs in a single bivalent IgG1 autoantibody to bind sites within the same receptor, consistent with previous findings^59,60^. However, the antibody orientation enables binding to an adjacent AChR and crosslinking of at least two receptors, given the high density of muscle AChR at the NMJ that is stabilized by the scaffolding protein rapsyn^61-64^. Superpositions of the receptors with the IgG modeled on the Fab binding site suggest that most of the autoantibodies studied here can lead to muscle receptor crosslinking, consistent with our previous functional results^12^ (Figures S8B–S8H, right-hand panels). We noticed that other epitopes remain available for polyclonal antibody binding on each receptor. However, whether crosslinking of two receptors is sufficient, or if larger-scale clustering is required to initiate internalization, requires further study. Interestingly, we found that the IgG1 structural constraints, with relatively short Fab-Fc hinge regions, would require membrane deformation to allow crosslinking (Figure S8). This inference is similar to a previous conclusion from a rat experimental MG antibody mAb35^59^. The Fab binding angle appears to be related to the membrane curvature; some mAbs would only require a slight distortion, while some would stabilize a large curvature coincident with crosslinking. Further research is needed to understand whether the membrane curvature favored by antibody-mediated crosslinking can trigger internalization, especially knowing that the endocytic process is effectively triggered by divalent MG antibodies^65^. In summary, while speculative, our structural analyses suggest that the pathologic effects of MG autoantibodies, such as complement activation and receptor internalization, are dependent on specific spatial binding angles and orientations.

In MG patients, polyclonal autoantibodies with different epitope specificities in activities may synergize to amplify pathogenic effects^7,12^. Antibodies that efficiently activate complement and/or trigger internalization likely depend on an optimal membrane organization of multiple antibody-receptor complexes, which is afforded by certain spatial orientations of the Fab with the receptor. In contrast, direct inhibition of receptor activity is less dependent upon these variables. Nevertheless, coincident interactions of diverse polyclonal antibodies in MG patient serum could either synergize or compete in mediating pathology. These interactions may explain the lack of correlation between antibody titers and disease severity.

### MG epitope landscape

Previous studies have concluded that many muscle receptor autoantibodies specifically target the α1 subunit, and specifically a region of the α1 subunit called the MIR^40,66,67^. This region is located on the upper lip of the α1 ECD, far away from the ACh site. This distance helps to explain the observation that MIR-directed autoantibodies generally do not directly affect receptor function^68^. Consistent with previous reports, we found that the α1 subunit harbors the most epitope determinants^41^. All MG Fabs, except for Fab9, interact with the α1 subunit. However, to our surprise, none of them directly contact the α1 MIR, revealing that many pathogenic autoantibodies in fact can target epitopes outside of the α1 MIR. Indeed, most patients have autoantibodies with multiple subunit specificities^7,12,69,70^. Interestingly, we found that one Fab could target the β1 MIR-like region and has an inhibitory effect on the channel function at a lower ACh concentration even though this site is also distant from the ACh site. The most common epitope regions in this study are on the outside surface of each ECD, which has a large surface area and is structurally accessible. Interestingly, some Fabs have similar binding positions but different patterns of epitope determinants. Our structures also show that the ECD vestibule inner surface can be targeted by autoantibodies. These findings further highlight the heterogeneity of binding position and epitope of AChR autoantibodies. The ε subunit has the fewest epitopes, which is consistent with the previously characterized specificity of monoclonal antibodies^7^. This rarity of ε epitope determinants is likely a result of the process of AChR-driven antibody maturation. Muscle-like myoid cells in the human thymus only express the fetal AChR isoform, which does not contain the ε subunit^71^. Structural analysis of additional monoclonal samples as well as polyclonal samples will provide a more comprehensive understanding of the MG antibody epitopes.

### Conclusion

In summary, our study provides key insights into the diverse pathological mechanisms underlying myasthenia gravis by examining the structure and function of the human muscle AChR bound to diverse MG autoantibodies. We defined distinct binding positions and mapped an epitope landscape across the muscle receptor, highlighting the complexity and heterogeneity of MG autoantibodies. These findings enhance our understanding of the mechanisms by which autoantibodies interfere with ion channel function and trigger indirect pathogenic processes, such as complement activation and receptor internalization. Importantly, this structural and functional knowledge can inform the development of more precise diagnostic tools and novel therapeutic strategies aimed at predicting and treating disease relapses. Moreover, these insights extend beyond MG, offering a broader perspective on the pathology of other antibody-mediated autoimmune disorders.

## Methods

### Engineering a cell line expressing the human adult muscle AChR

We first tested recombinant expression of the adult human muscle ACh receptors by transiently transfecting HEK293 cells with four plasmids each containing the individual subunit genes (α1, β1, ε, δ). To evaluate the assembly of these subunits into heteropentamers of correct composition, versions of these plasmids were used that either did or did not contain GFP fused into their intracellular M3M4 loops. Cells were transfected with all single, double, triple and quadruple combinations of plasmids wherein only one subunit was tagged with GFP, to define rules of assembly in this overexpression system. Using a fluorescence-detection size-exclusion chromatography (FSEC)^72^, we found that the α1, β1 and ε subunits individually can form apparent homopentamers, but that the δ subunit cannot and instead is only incorporated into heteropentamers. This result suggested that the δ subunit was suitable for affinity tag fusion. To streamline protein expression, we aimed to set up a stable cell line, to avoid cumbersome large-scale transfections or transductions with four plasmids or viruses. The Sleeping Beauty system^73^ is a rapid method for generating stable cell lines, but is limited in the size of inserted genes^74^. Accordingly, we separated these four genes into two pairs. To minimize expression of incorrect subunit assemblies, we tested different subunit combinations. Again, for each test, only one subunit contained a GFP fusion. We found that paired combinations of α-δ and β-ε genes generated mostly correct heteropentamers. We thus subcloned these gene pairs into two different Sleeping Beauty plasmids, each of which contained different selection and fluorescence markers. Each pair contains two full-length wild-type genes (without GFP) and that are linked by a T2A self-cleaving peptide gene. The M3M4 loops of the δ and ε subunits include inserted Twin-Strep and 3XFLAG tags respectively for affinity purification.

To generate the stable cell line, 1 μg pSBtet-GP-α+δ, 1 μg pSBtet-RH-β+ε and 0.1 μg pCMV(CAT)T7-SB100 plasmids were transfected into HEK293S GnTI^-^ cells (ATCC CRL-3022) at 70% confluency using Lipofectamine 2000 (Invitrogen). The two pSB Sleeping Beauty system vectors were a gift from Eric Kowarz and obtained from Addgene (Catalog #60495, 60500)^73^ and the SB transposase plasmid was a gift from Zsuzsanna Izsvak, also obtained from Addgene (Catalog #34879)^75^. After 24 hours, 1.5 μg/mL puromycin and 100 μg/mL hygromycin B were added for positive cell selection. The dual selection was maintained for two weeks until all transfected cells exhibited both GFP and dTomato fluorescence. Cells were next split into 10 cm dishes and then into 2 L suspension flasks, in FreeStyle 293 medium (ThermoFisher) + 2% fetal bovine serum (Sigma), for large-scale protein production.

### Optimization of expression and purification of human muscle AChR

We screened different culture temperatures, concentrations of induction reagent, expression boosters, and time-courses of expression. Briefly, stably-expressing cells were plated into a 6-well dish and cultured at 37 °C. Different tests were performed using these dishes once cells reached 70% confluency. We found that protein expression is higher at 30 °C compared to 37 °C, after induction. One μg/mL doxycycline was optimal for induction. Expression boosters like sodium butyrate did not noticeably improve receptor expression. The peak expression occurred 2 days after induction.

For purification, we tested solubilization using different detergents and cryo-EM analysis on the large-scale suspension culture. Three to 5 L cell cultures were collected and membrane pellets were prepared using previous protocols^76^. The membranes were solubilized in buffers containing 40 mM DDM (*n*-dodecyl-β-D-maltoside, Anatrace), or 10 mM LMNG (lauryl maltose neopentyl glycol, Anatrace), or 2% Triton X-100 (Sigma), or 20 mM DDM + 0.5% GDN (glyco-diosgenin, Anatrace), or 1% GDN, and then exchanged to corresponding detergents. Soy lipids and cholesterol were added throughout the test purifications. In cryo-EM analysis, we found that the TMD and ICD were sensitive to the extraction conditions, wherein most solubilizations led to a partially damaged TMD and disordered ICD. By contrast, solubilization in 1% GDN was suitable for human muscle AChR and generated a stable and homogeneous protein sample.

### Human muscle AChR preparation for high-resolution single-particle analysis

5 L suspension cultures of stably-expressing cells were harvested by centrifugation at 5,000 × g, 15 min, 4 °C, after 2 days of induction at 30 °C. Cell pellets were resuspended in a buffer containing 50 mM Tris-HCl pH 7.4, 150 mM NaCl, 2 mM EDTA, cOmplete protease inhibitor cocktail (Sigma) and 1 mM PMSF (phenylmethylsulfonyl fluoride). The cell suspension was then passed four times through an Avestin C5 cell disruptor at 5,000-10,000 psi. The lysate supernatant was collected after centrifugation at 10,000 × g, 15 min, 4 °C and re-centrifuged at 40,000 rpm, 4 °C for 2 hours using a 45Ti rotor to pellet the cell membranes. Membranes were stored at -80 °C until needed.

For each purification, approximately 10 g of membranes were Dounce homogenized in a buffer containing 50 mM Tris-HCl pH 7.4, 150 mM NaCl, 1 mM EDTA, 1 mM PMSF, cOmplete protease inhibitor cocktail (Sigma) and 10 µM soy polar lipids (Avanti Polar Lipids) plus cholesterol (Sigma) and DDM in a 4:1:1.84 w:w ratio). GDN was then added to a final concentration of 1%, and solubilization was performed at 4 °C while nutating for 3 hours. Insoluble material was removed by centrifugation at 186,000 g for 40 min. The supernatant containing target proteins was bound to 3 mL of Strep-Tactin XT 4Flow high-capacity resin (IBA Life Sciences) using a peristaltic pump at 0.8 mL/min, 4 °C. The resin was washed with 200 mL TBS plus 0.5 mM GDN supplemented with 10 µM soy polar lipids plus cholesterol to stabilize the receptors. The receptor proteins were eluted in the same TBS buffer with 10 µM soy polar lipids plus cholesterol and 0.25 mM GDN supplemented with 50 mM biotin (Sigma). Elution samples were analyzed by FSEC using intrinsic tryptophan signal, pooled, and concentrated for preparative SEC (in 50 mM Tris-HCl pH 7.4, 150 mM NaCl, 0.25 mM GDN, and 10 µM soy lipids plus cholesterol) using a Superose 6 Increase 10/300 GL column (Cytiva) and SDS-PAGE analysis. For the ACh-bound receptor purification, the SEC buffer was supplemented with 1 mM ACh. Peak SEC fractions were pooled and prepared for cryo-EM or Fab complex formation.

### Fab fragment preparation

The heavy and light chain variable region DNA sequences of MG patient mAbs 1b, 2, 3, 6, 7, 9 were used for the generation of monovalent Fabs. DNAs synthesized by GenScript were tagged with 8xHIS and 3XFLAG sequences at their heavy chain C termini and subcloned into the pCEP4 vector (Invitrogen) for high-level, constitutive protein expression in mammalian cells. These endotoxin-free Fab plasmids were purified using the PureLink HiPure Plasmid Maxiprep Kit (Invitrogen, catalog: K210007) and were transfected into ExpiCHO cells (Thermo Fisher). For large-scale expression, 200 µg of each heavy and light chain plasmid were transfected into 400 mL cells at a density of 6 x 10^6^ cells per mL using the ExpiFectamine CHO Transfection Kit (ThermoFisher, catalog: A29129). After 20 hours of transfection, the Enhancer (2.4 mL) and Feed (96 mL) reagents were added into the cell culture, which generated approximately 500 mL of cell suspension for each Fab preparation. After 8 days in culture, cells were harvested by centrifugation at 5,000 × g, 30 min, 4 °C. The supernatant was loaded onto a His Trap HP column (Cytiva) using a peristaltic pump at 1.6 mL/min. To ensure sufficient binding, the flow-through was recirculated through the column for 18 hours at 4 °C. The column was washed using a buffer (50 mM Tris-HCl pH 7.4, 150 mM NaCl, 25 mM imidazole) to remove nonspecific binding proteins. Fabs were eluted with the same buffer but containing 500 mM imidazole, and were further polished using a Superdex 200 Increase 10/300 GL column (Cytiva). The peak fractions were pooled and analyzed by SDS-PAGE. The purified Fabs fragments were stored at -80 °C for electrophysiology and cryo-EM analysis.

### Preparation of AChR-Fab complexes

For Fab3, 6, 7 and 9, the purified muscle AChR proteins were mixed with Fabs at a molar ratio of 1:2 and incubated for 40 mins on ice. The mixtures were concentrated and further polished using a Superose 6 SEC column (Cytiva) with a mobile phase containing 50 mM Tris-HCl pH 7.4, 150 mM NaCl, 0.25 mM GDN and 10 µM soy lipids plus cholesterol, and the peak fractions were pooled and concentrated for further analysis and cryo-EM sample preparation. For Fab1b and Fab2, the purified muscle AChR proteins were mixed with Fabs at a molar ratio of 1:5 and incubated for 40 mins on ice. The excess Fabs were removed using 100 kD MW cutoff concentrators by adding 4 mL buffer (50 mM Tris-HCl pH 7.4, 150 mM NaCl, 0.25 mM GDN and 10 µM soy lipids plus cholesterol). The samples were concentrated for further cryo-EM analysis.

### Cryo-EM sample preparation

Three μL of freshly purified muscle AChR proteins or the polished Fabs complexes at a concentration of 0.2 mg/mL were added to a freshly glow-discharged (at 30 mA for 10 s) 200-mesh copper grid containing a 2 nm carbon film (R2/1, Quantifoil). Grids were waited for 20 s and blotted for 3.5 s under 100% humidity at 4 °C with blot force -15. Grids were plunge-frozen into liquid ethane using a Vitrobot. Grid quality was checked on the UCSD 200 kV Talos Arctica microscope. The grids with good ice, particle distribution, and density were saved.

### Cryo-EM data collection and image processing

The data processing strategies for apo and ACh-bound datasets were similar as were all Fab complex datasets. Here we cover the processing details for apo and Fab6-bound datasets, as two representative examples. For the apo dataset, 8,004 raw images were collected on the UCSD Titan Krios 2 at 300 kV with a total dose of 50 e^-^/Å^2^ and a magnification of 130,000x, resulting in a pixel size at 0.935 Å/pixel. The defocus range was set from -1.4 to -2.0 µm for the collection. All data processing was done using cryoSPARC v4.4^77^. The gain-normalization, motion correction, and CTF estimation were done using default parameters in cryoSPARC. Approximately 6.2 million particles were picked using the blob picker and extracted with a box size of 400 pixels and binned to 200 pixels. After five rounds of 2D classification, approximately 2 million good particles were selected from classes with clear secondary structural features. Twenty thousand particles were randomly selected to generate a 3D model using *ab initio* reconstruction. All particles were then submitted into a 3D Homogeneous Refinement job using the *ab initio* model as the initial volume. To improve particle alignment during 3D classification, two soft focus masks were generated using Relion_mask_creation^78^ based on the 3D homogeneous refinement map. First, the mask was focused on the ECD to remove particles with poor local ECD densities such as damaged loops and glycosylation using 3D classification (10 classes), then all good particles with complete ECDs were selected. To improve TMD and ICD density, the second focus mask on TMD and ICD was applied for the second 3D classification (10 classes). This step separated the complete particles (1,128,019 particles) from those without ICD density, and weak TMDs. In both 3D classifications, the mode was set to PCA and the convergence criterion was changed to 1%. To further remove the remaining bad particles, an *ab initio* reconstruction in combination of Heterogeneous Refinement were performed, which generated only one class with all parts and high quality. This class was selected, re-extracted at full size, increased box size to 512 pixels and refined using cryoSPARC NU Refinement^79^ to a resolution of 2.05 Å (556,873 particles).

The Fab-bound samples’ EM data were comparatively straightforward to process because Fab binding facilitated particle alignment. For the Fab6-bound sample, 7,697 raw images were collected using the settings as above. All data processing was again performed using cryoSPARC v4.4. After four rounds of 2D classification, approximately 1.3 million particles were kept and subjected to a 3D Homogeneous Refinement using an initial model generated from an *ab initio* reconstruction. The 3D Homogeneous Refinement result was then submitted to a 3D classification (8 classes) with PCA mode and 1% convergence criterion. This provided three classes with clear Fab binding and strong TMD and ICD densities. These classes were kept and further subjected to an *ab initio* reconstruction in combination with Heterogeneous Refinement, which removed few bad particles. Only one intact receptor class was selected, re-extracted at full size, and refined using cryoSPARC NU Refinement. To further improve the resolution, the local CTF refinement and Reference Based Motion Correction were performed. The final NU Refinement using the particles after polishing generated a map at a resolution of 1.92 Å (543,540 particles).

### Model building, refinement, and validation

Given the high quality and resolution of cryo-EM maps, we used ModelAngelo^80^ for initial *de novo* model building. This auto building software generated one model that fitted the apo muscle AChR map very well using supplied sequences. This model, which included several fragments, was linked and checked manually residue by residue in Coot (version 0.9.8.93)^81^ and refined using global real space refinement in Phenix (version 1.20.1)^82^ with secondary structure and Ramachandran restraints. Model geometry and clash scores were checked using MolProbity^83^. For the ACh-bound map, the well-refined apo structure was used as the starting model, and roughly fitted into the new density map using UCSF Chimera (version 1.16)^84^. The most different regions including the channel pore and some loops fitted poorly and were firstly manually rebuilt, and then subjected to several cycles of real space refinement in Phenix. ACh molecules were positioned into unambiguous density in the classical neurotransmitter sites. Finally, the improved model was then checked manually residue by residue in Coot and in combination with several rounds of global real space refinement in Phenix with secondary structure and Ramachandran restraints. Model geometry and clash scores again were checked with MolProbity.

For Fab-bound maps, the structures of Fab molecules were also *de novo* built by ModelAngelo with the Fab sequences provided. For the constant region, which has a lower local resolution, ModelAngelo could not build a complete backbone; we thus fitted a high-resolution crystal structure of the constant region of human IgG (PDB 4LLD)^85^ into these regions using UCSF Chimera and refined the model by Phenix. For the receptor region, the well-refined apo structure again was used as the starting model, and roughly fitted into the new density maps and then refined by Phenix in combination with Coot. Model geometry and clash scores were checked with MolProbity.

All model building was performed based on the final sharpened maps from cryoSPARC NU Refinement while DeepEMhancer^86^ was only used to improve the map quality of some regions with lower local resolution. Sequences for human muscle receptor model building were downloaded from the UniProt database and sequence alignments were made using Clustal Omega^87^. The pore diameter was calculated using HOLE2^88^. The map and structural figures were generated using UCSF Chimera, ChimeraX (version 1.7.1)^89^ and PyMOL (version 2.5.5, Schrodinger, LLC). The main text figures were made from density maps after post processing with DeepEMhancer. Some lipids densities were not strong enough, but were built based on consistency with our earlier bovine muscle AChR models^15^.

### Mass spectrometry analysis

Proteins were identified using mass spectrometry in the UCSD proteomics facility after SDS-PAGE separation based on standard methods^90^. Protein bands around 50 kD were cut from a Coomassie blue-stained gel using a sterile blade. The gel slices were diced into 1 mm³ cubes and subjected to destaining with 100 mM ammonium bicarbonate and acetonitrile. Following destaining, the sample was reduced, alkylated, dehydrated, then were processed with trypsin digestion overnight. Post-digestion, the peptides were extracted and analyzed by liquid chromatography-tandem mass spectrometry (LC-MS/MS) with electrospray ionization. The analysis was performed using the Bruker TimsTOF 2 pro mass spectrometer in conjunction with a nano-scale reversed-phase UPLC system (EVOSEP ONE). Mass spectrometry was conducted with a PASEF method incorporating mobility-dependent collision energy ramping and precursor target intensity adjustments. Protein identification and label-free quantification were performed using Peaks Studio X software.

### Electrophysiology

Whole cell voltage-clamp recordings were made from adherent HEK293S GnTI^−^ cells stably expressing doxycycline inducible human α1, β1, δ, and ε subunits (see above for details on the stable inducible expression system) or transiently transfected with human α1, β1, δ, and ε. After 48 hours transfection or induction by 1 µg/mL of doxycycline, cells were re-plated on 35 mm dishes and allowed to settle for at least 3 hours.

Recordings were made 48-96 hours after induction. The bath solution contained (in mM): 140 NaCl, 2.4 KCl, 4 MgCl_2_, 4 CaCl_2_, 5 HEPES and 10 glucose pH 7.3. Borosilicate pipettes were pulled and polished to an initial resistance of 2–4 MΩ, filled with the pipette solution containing (in mM): 100 CsCl, 30 CsF, 10 NaCl, 10 EGTA, and 20 HEPES pH 7.3. Whole cell currents were recorded with an Axopatch 200B amplifier, sampled at 20 kHz, and low-pass filtered at 2 kHz using a Digidata 1550B (Molecular Devices). Cells were held at -75 mV. Solution exchange was achieved using a gravity driven RSC-200 rapid solution changer (Bio-Logic). Whole cell currents data were analyzed with Clampfit 11 software (Molecular Devices). 2 or 300 µM of Acetylcholine and 10 µg/mL of Fabs were prepared in bath solution from concentrated stocks, 1 M ACh in water, and 1∼10 mg/mL Fabs in PBS stored at -80 °C.

### Generation of mutant AChR-specific recombinant autoantibodies

Mutants of native AChR-specific mAbs were designed by mutating select amino acid residues to either an alanine if the residue was not generated by SHM, or its germline-encoded amino acid residue, which was assigned through V-gene alignment using the IMGT (IMmunoGeneTics) V-Quest database^37^. Mutant mAb expression plasmids were generated by DNA fragment synthesis of each heavy and/or light chain variable region (Twist Biosciences) which were cloned into human IgG1 or Igκ expression vectors^91^. Mutant plasmids were verified via Sanger sequencing with a CMV forward primer located upstream of the antibody V(D)J. Recombinant autoantibodies were expressed by transfection of heavy and light chain plasmid pairs into HEK293T cells (ATCC, CRL3216) in 6-well plates (Falcon, 353046) in the presence of linear polyethylenimine (Polysciences, 23966) in complete DMEM (Dulbecco’s Modified Eagle Medium) (Gibco, 11995) supplemented with 10% heat-inactivated fetal bovine serum (Sigma, F0926), 1% L-glutamine (ATCC, 30-2214), 1% Non-essential amino acids (Gibco, 11140050), and 1% Penicillin-Streptomycin (Gibco, 15140122). Cells were incubated at 37 °C, 5% CO_2_ for 16 hours after which the culture medium was removed and replaced with serum-free basal medium supplemented with 1% Nutridoma-SP (Roche, 11011375001). After 3 additional days of incubation, antibody-rich culture supernatants were harvested from HEK293T cells.

### Recombinant antibody quantification by ELISA

Recombinant autoantibody concentrations in cell culture supernatants were measured by ELISA. ELISA plates (96-well) were coated overnight at 4 °C with 5 μg/mL of Fcγ fragment specific, goat anti-human IgG (Jackson ImmunoResearch, 109-005-098) and subsequently blocked for 1 hour at 23 °C in blocking buffer (1% BSA, 0.05% Tween-20 in PBS). Antibody-containing supernatant were then added to the plates and incubated for 1 hour at 23 °C, then washed, followed by addition of 0.04 μg/mL secondary anti-IgG-HRP antibody (Jackson ImmunoResearch, 109-036-098). The ELISA was developed with TMB ELISA substrate solution (ThermoFisher, 34028) for 2 minutes, acidified with 1 M H_2_SO_4_, and immediately measured with a spectrophotometer at 450 nm. Absorbance values were converted to concentrations (μg/mL) using a standard curve established with a human IgG standard (Sigma, I4506).

### Rapsyn-clustered cell-based assay (CBA) for assessing AChR autoantibody binding

AChR-specific autoantibody binding was measured using a live cell-based assay. HEK293T cells were transfected with human adult AChR (2α1, β1, δ, ε) and rapsyn-GFP plasmids (generous gifts from Drs. Angela Vincent, David Beeson, and Patrick Waters, Neurosciences Group at the Weatherall Institute of Molecular Medicine, University of Oxford) in the presence of branched polyethylenimine (Sigma, 408727). Culture medium was replenished the next day, and cells were harvested for autoantibody testing the following day. The resulting HEK293T cells were then seeded into 96-well format at 50,000 cells per well and stained with recombinant AChR antibody-containing supernatants 4-fold serially diluted from 1 to 0.02 μg/mL for 1 hour at 4 °C. Cells were washed twice, and subsequently stained with a secondary anti-human-IgG Fcγ Alexa-Fluor™-647-conjugated antibody (Jackson ImmunoResearch, 309-605-008) to detect primary autoantibody binding. Positive AChR-specific binding on cells was determined by gating on AChR/Rapsyn^+^ IgG^+^ cells on an LSR Fortessa flow cytometer (BD Biosciences). Flow cytometric data was analyzed using FlowJo software (v10.10.0). AChR-specific autoantibody binding capacity was quantified by a normalized mean fluorescence intensity (ΔMFI) calculated as (ΔMFI = MFI_AChR-GFP-positive_ - MFI_AChR-GFP-negative_). An anti-aquaporin-4 antibody (mab58)^92^ (a generous gift from Dr. Jeffrey L. Bennett of the University of Colorado) was included in each assay as a non-AChR binding IgG negative control.

### Quantification and statistical analysis

The main text figures describing electrophysiology results are presented as normalized peak currents.

Replicate numbers from independent cells are labeled in each bar. Statistical analysis was performed using GraphPad Prism 10.2.0 software (GraphPad software, Inc, La Jolla, CA). The two-tailed Welch’s t-test was used for electrophysiology. Statistical testing for all CBAs was performed using the two-way ANOVA with Dunnet’s correction. The *P*-values are shown in the related Figures. Data are expressed as mean ± s.e.m.

## Data availability

All atomic models and cryo-EM maps have been deposited in the Protein Data Bank and Electron Microscopy Data Bank: Apo/resting (PDB 9DMG, EMD-47003); ACh-bound/desensitized (PDB 9DMH, EMD-47005); Fab1b-bound, two Fabs (PDB 9DMJ, EMD-47007); Fab1b-bound, one Fab (PDB 9DMK, EMD-47008); Fab2-bound (PDB 9DML, EMD-47009); Fab3-bound (PDB 9DMQ, EMD-47013); Fab6-bound (PDB 9DMS, EMD-47014); Fab7-bound (PDB 9DMT, EMD-47015) and Fab9-bound (PDB 9DMV, EMD-47017).

## Acknowledgements

We thank S. Burke, W. Chojnacka, J. Zhou, and H. Jiang for critical feedback on the manuscript and L. Baxter for assistance with figures. We also thank the UC San Diego Cryo-EM Facility and its staff members, M. Matyszewski and I. Kuschnerus, for their scientific and technical assistance; M. Ghassemian for mass spectrometry support. MCP was supported by an NIH training grant (AI007019). KCO was supported by grants from the NIH (AI114780 and AI164590). REH was supported by the Myasthenia Gravis Foundation of America and the NIH (NS120496 and NS130831).

## Author contributions

H. Li performed the biochemistry and structural biology experiments and drafted and revised the manuscript. M. Pham performed mAb mutagenesis, flow cytometry and binding assays and revised the manuscript. J. Teng performed the electrophysiology experiments. K. O’Connor oversaw the mAb experiments and revised the manuscript. C. Noviello conceived the project with R. Hibbs and drafted and revised the manuscript.

## Supplemental figures

**Figure S1.**
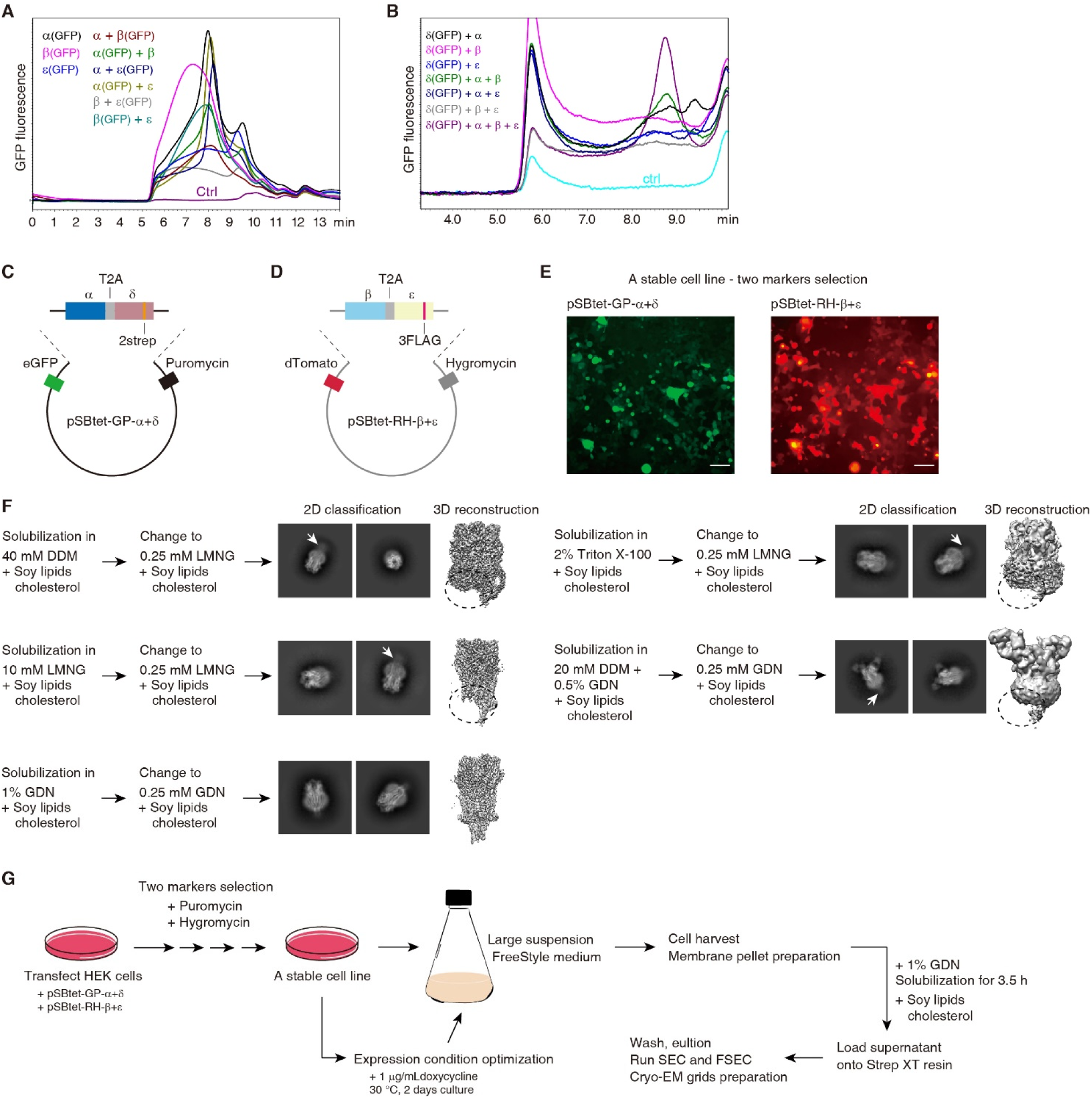
Design and optimization of human muscle AChR expression and purification. (A) Fluorescence-detection size-exclusion chromatography traces of individual muscle-type AChR subunits tagged with GFP or in combination with each other. X axis is time of elution; Y axis is GFP fluorescence. (B) Examination of receptor expression after transfecting different combinations of subunits, with only the δ subunit tagged with GFP. (C), (D) Construct design for stable-cell line generation. (E) Fluorescence from HEK cells transfected with plasmids from C, D indicates both constructs are integrated into the cell line. Scale bar, 200 μm. (F) Detergent optimization drastically improved ability to determine the TMD and ICD substructures. (G) Overview of final expression and purification strategy.

**Figure S2.**
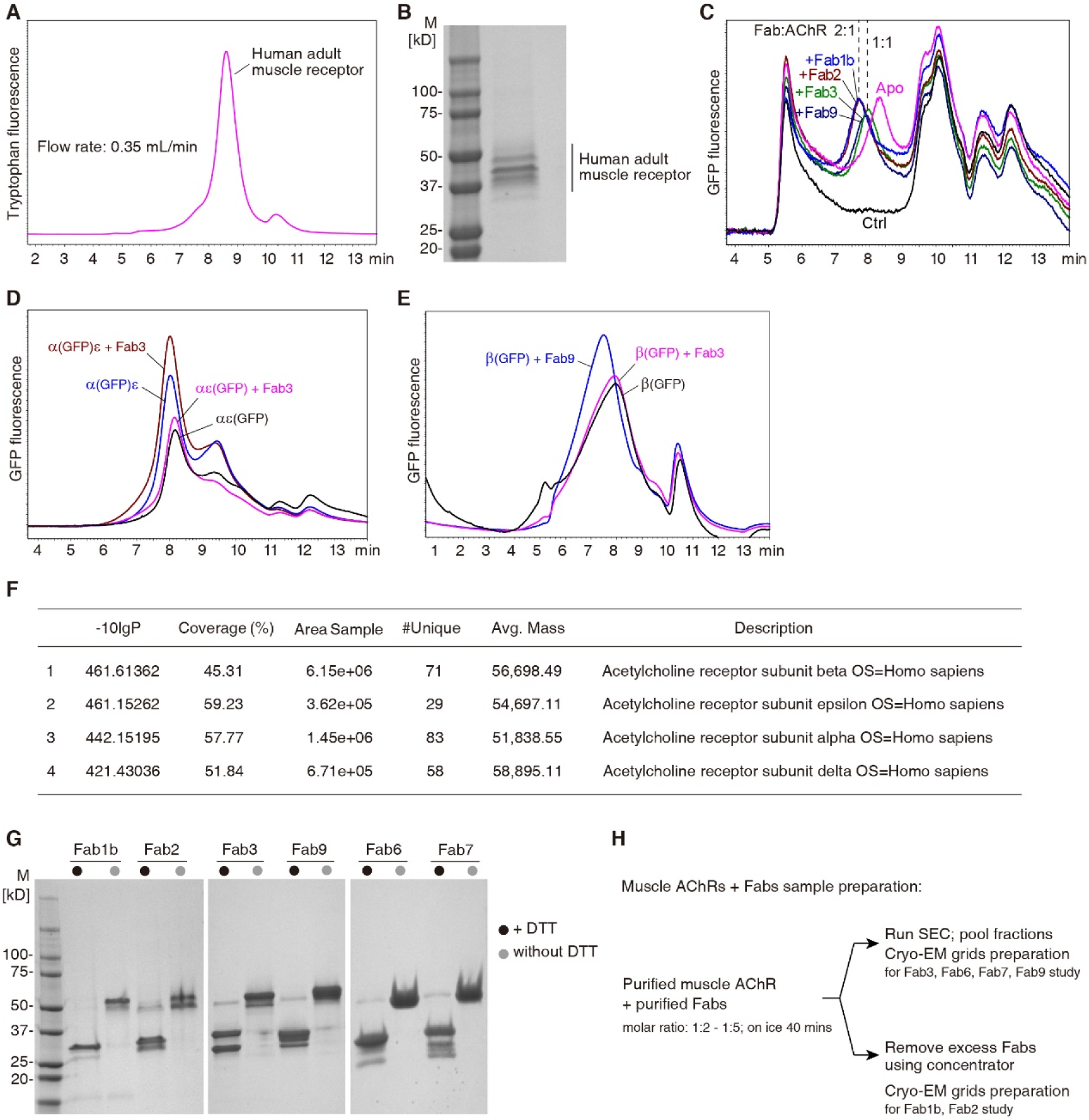
Expression and formation of muscle AChR-antibody fragment complexes. (A) FSEC trace of human adult muscle AChR as purified via workflow in Figure S1. Sepax SRT SEC 500 column with flow rate: 0.35 mL/min. (B) Coomassie staining of SDS-PAGE gel indicates subunits migrate at appropriate size. (C) FSEC binding tests of Fabs to receptor show binding via shift to different elution times, suggesting various stoichiometries indicated by dashed lines. (D), (E) FSEC binding tests of different Fabs show differing subunit specificity. (F) Mass spectrometry results from gel seen in panel B show presence of all four adult AChR subunits. (G) SDS-PAGE stained with Coomassie of overexpressed and purified Fabs used in this study. (H) Overview of purification and formation of receptor-antibody complexes for structure determination.

**Figure S3.**
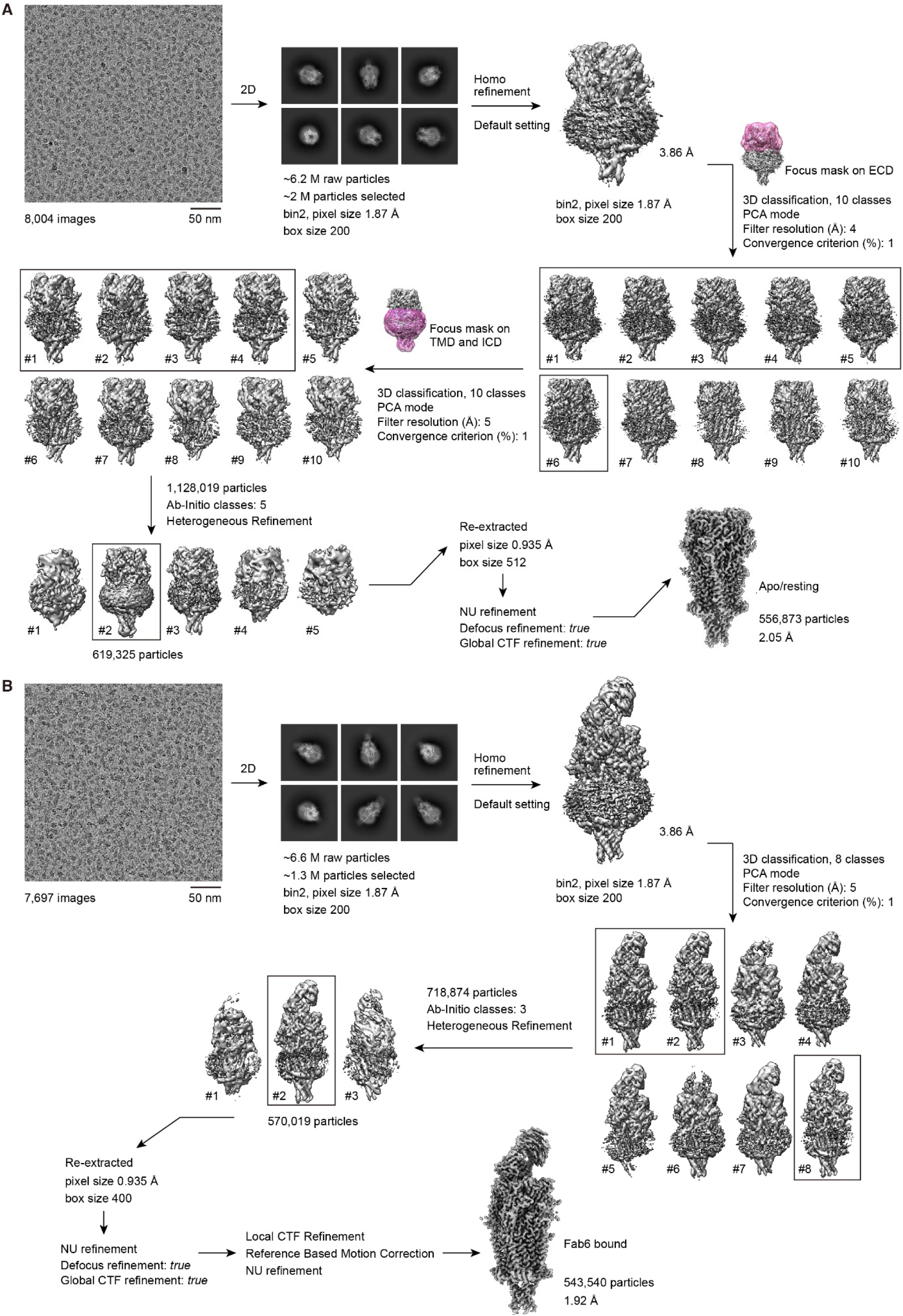
Representative cryo-EM data processing workflows. Representative cryo-electron microscopy workflow used to determine high-resolution structures of reference apo-resting (A) and Fab6-bound (B) AChR complexes. Each step is delineated by arrows and performed in CryoSPARC v4.4.

**Figure S4.**
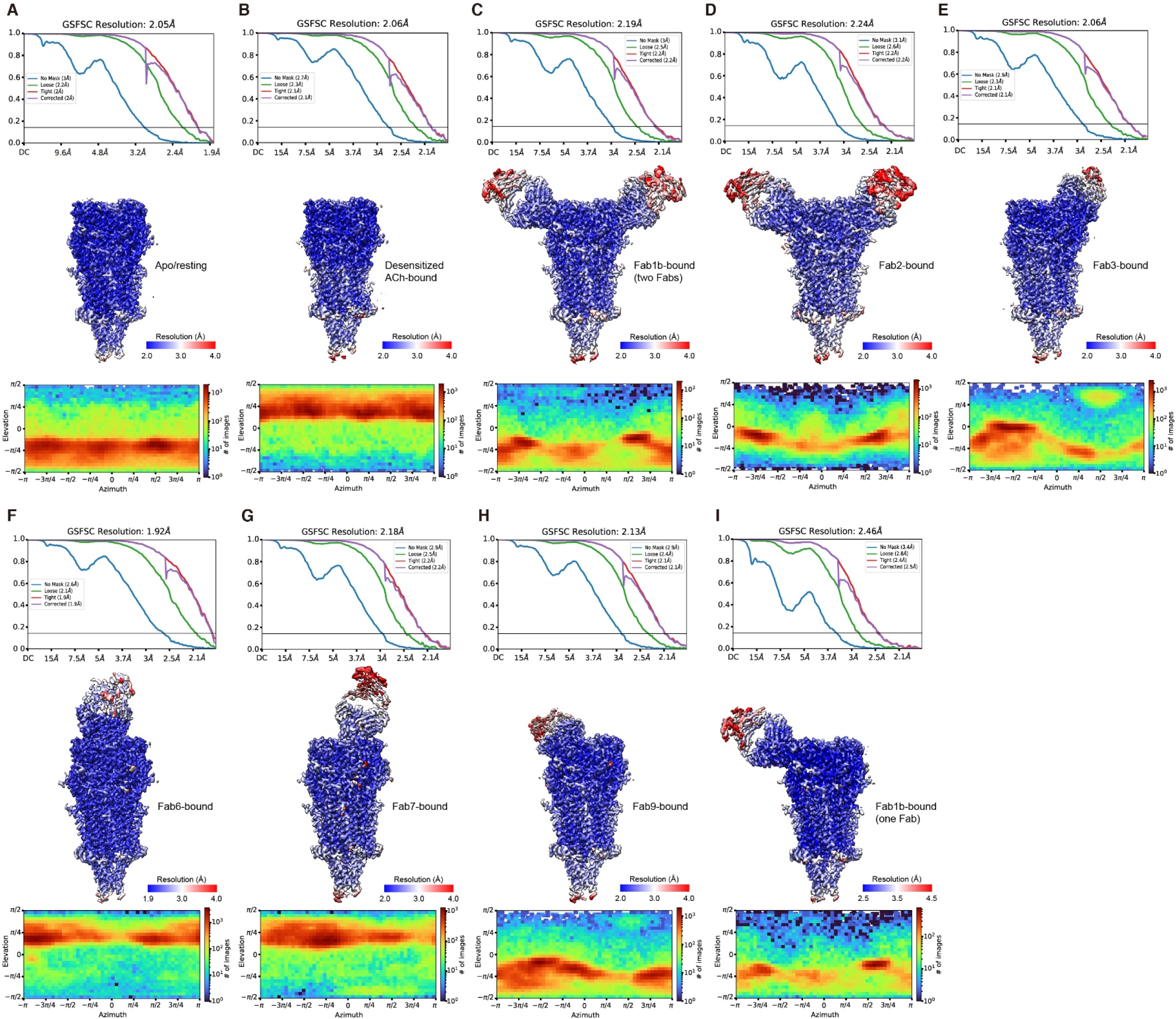
Map quality evaluation for the human muscle AChR and its antibody complexes. Each panel is for a different density map; top are Fourier-shell correlations, with resolution determined at gold-standard cut-off of 0.143 (solid line across graph). Unmasked, loose, tight, and corrected masks are plotted, with resolution of each indicated in parentheses. Maps beneath are color-coded for local resolution as indicated in scale bar on right. The bottom graphs of each panel are angular distribution plots.

**Figure S5.**
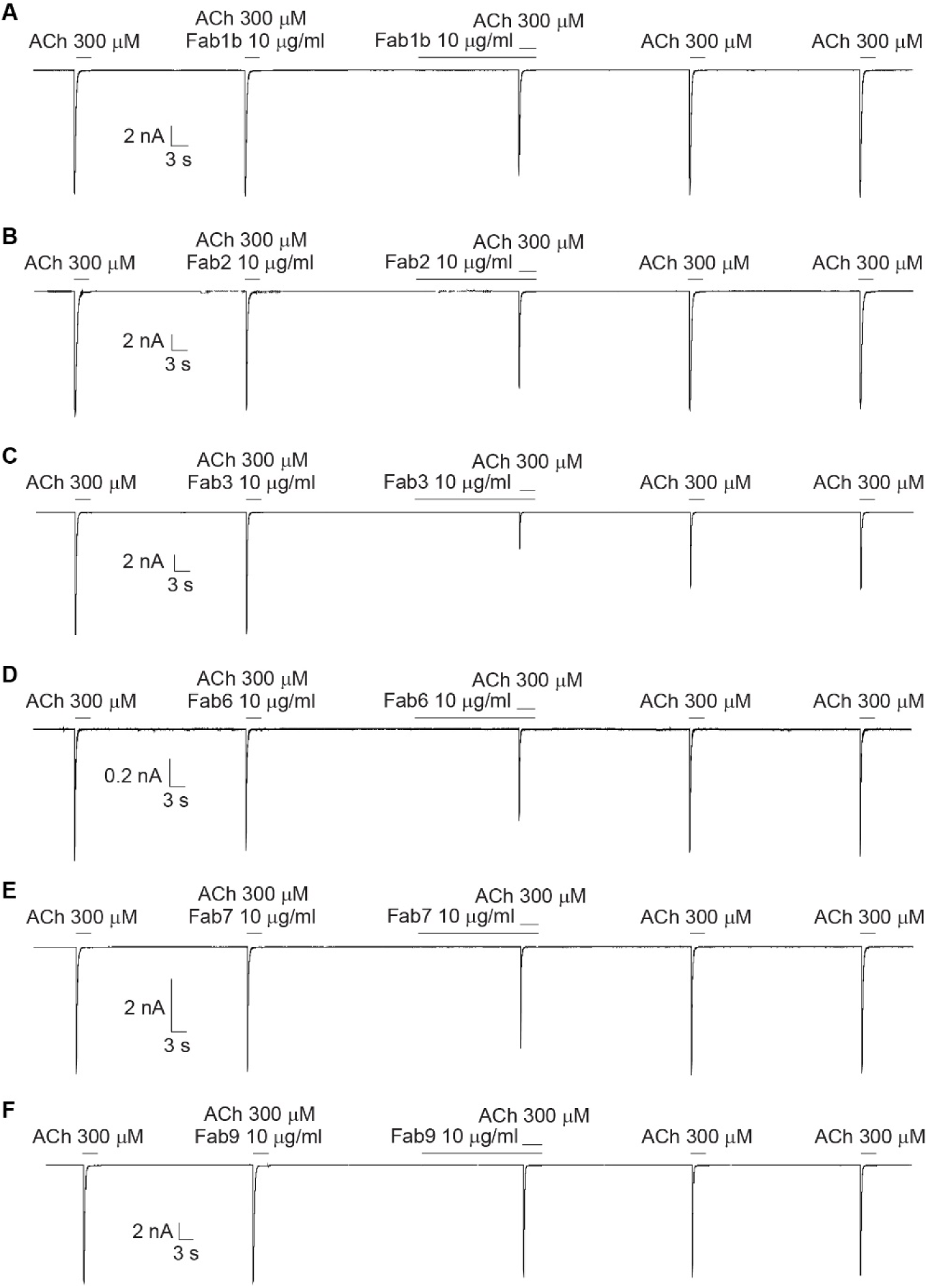
Electrophysiology tests of receptor inhibitions by different Fabs at high ACh concentrations. Representative raw traces of channel responses in the presence of different Fabs at 300 μM ACh in whole-cell patch clamp electrophysiology. Related to Figures 3, 4, 5 and 6.

**Figure S6.**
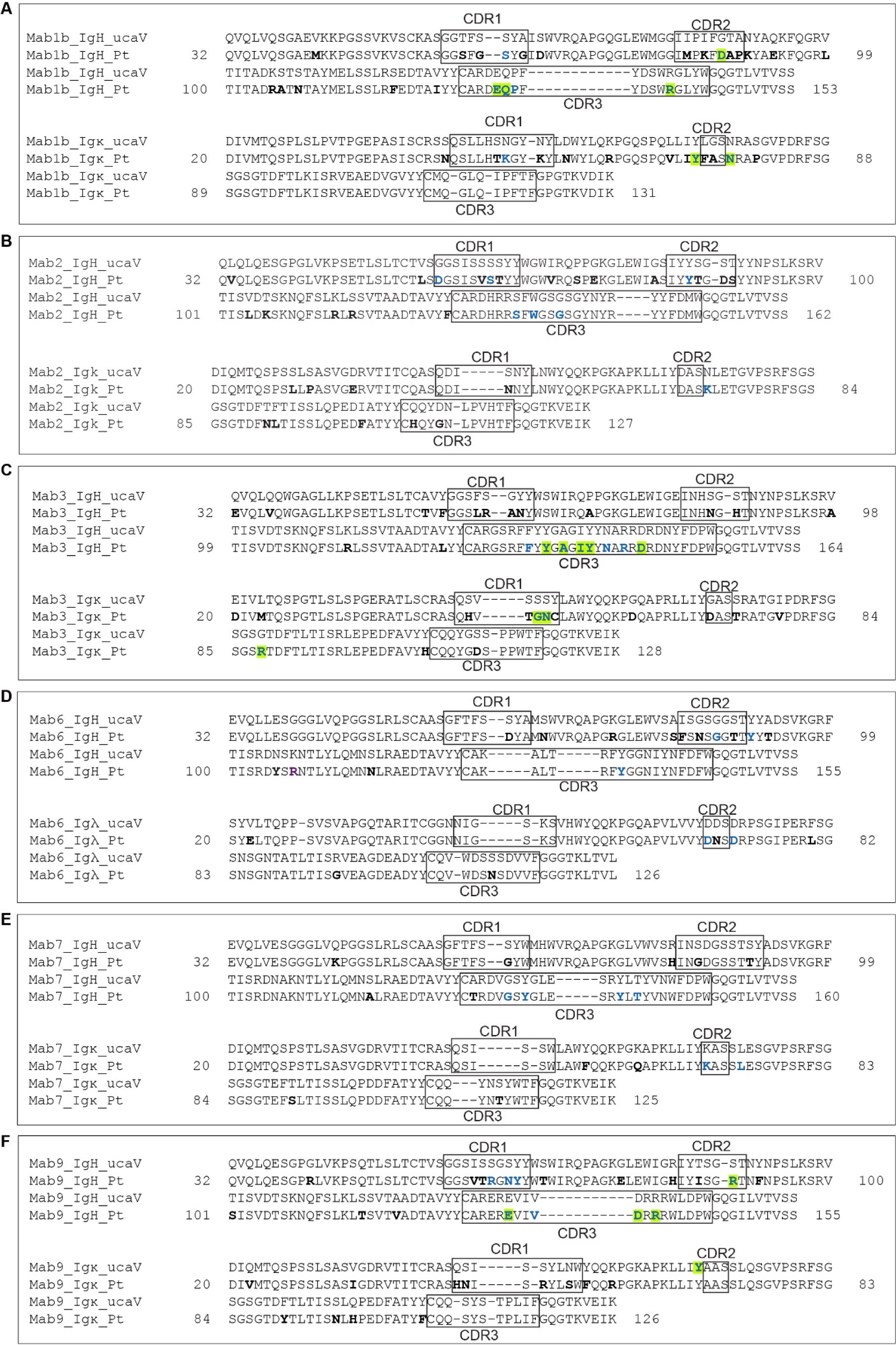
Patient antibody sequences aligned to unmutated common ancestor (germline) sequences. Variable domains of antibody sequences from patient-derived autoantibodies used in this study are shown aligned to the germline (ucaV) sequences of the variable gene. The diversity and junction segments are too variable to enable confidence in assignment of mutated residues versus germline sequences, and thus are left the same. Residues that are a result of somatic hypermutation are shown in bold. Residues that contact the receptor are indicated in blue. Residues selected for binding analysis are boxed in green. The complementarity determining regions (CDR) of each chain are indicated.

**Figure S7.**
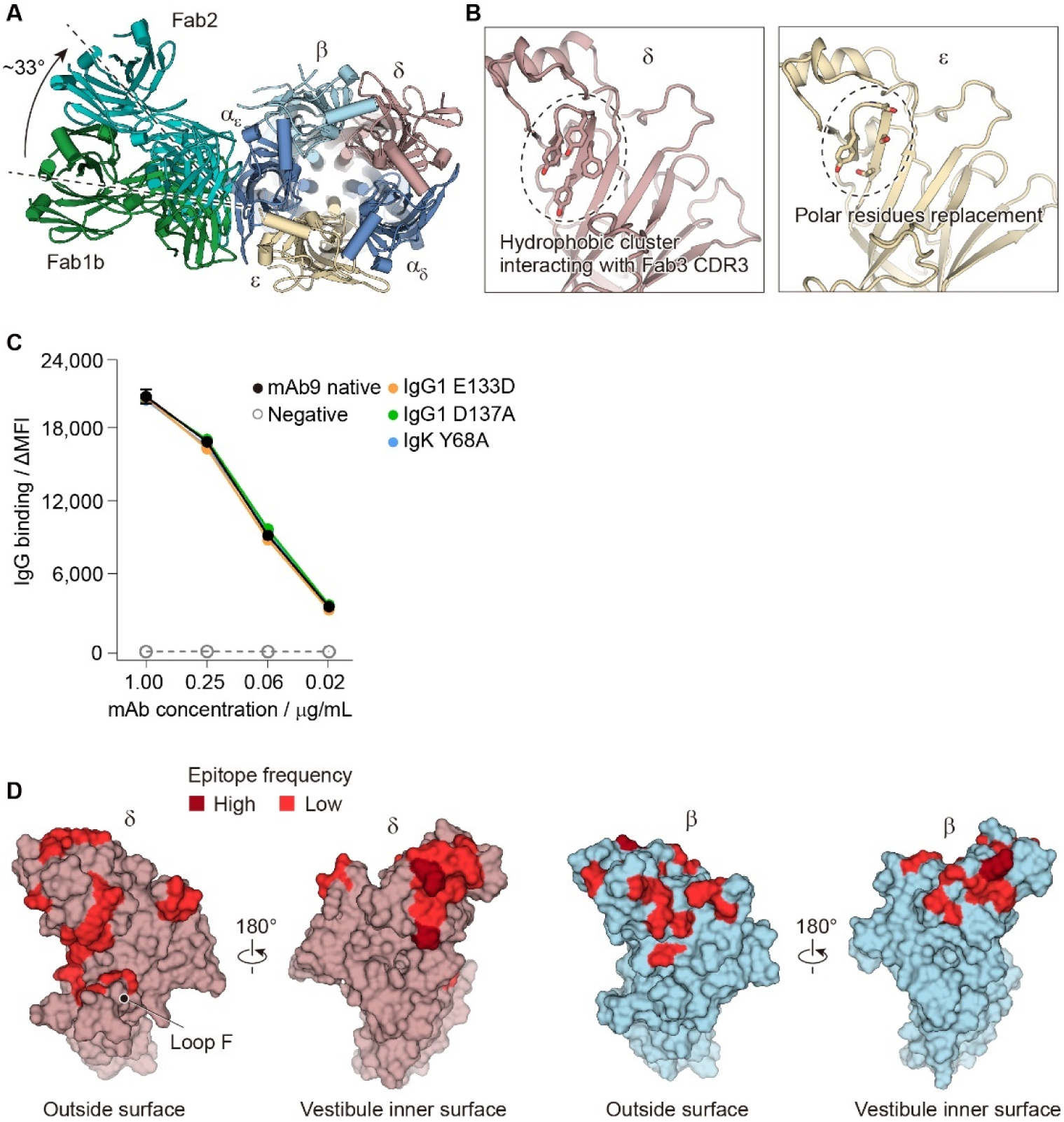
Structural and functional analyses for Fab-receptor structures and epitope distributions in the ECD vestibule. (A) Structural comparison of Fab1b- and Fab2-bound AChR structures reveals that Fab2 binds to a similar region of the α1 subunit but is rotated clockwise relative to Fab1b. (B) The hydrophobic cluster on the δ subunit interacting with the Fab3 CDR3 has been replaced by small polar residues in the ε subunit. (C) Titration plot binding curves of the native mAb9 and its mutants tested for binding over a range of serially diluted concentrations. Negative control, mAb58 (aquaporin-4 specific). Each data point represents a mean of experimental triplicate values. Data are expressed as mean ± s.e.m. Related to Figure 6. (D) Surface representations show epitopes in the ECD outside surface and vestibule inner surface of δ and β subunits. Color-coded by antibody interactions as determined by cryo-EM. Related to Figure 7.

**Figure S8.**
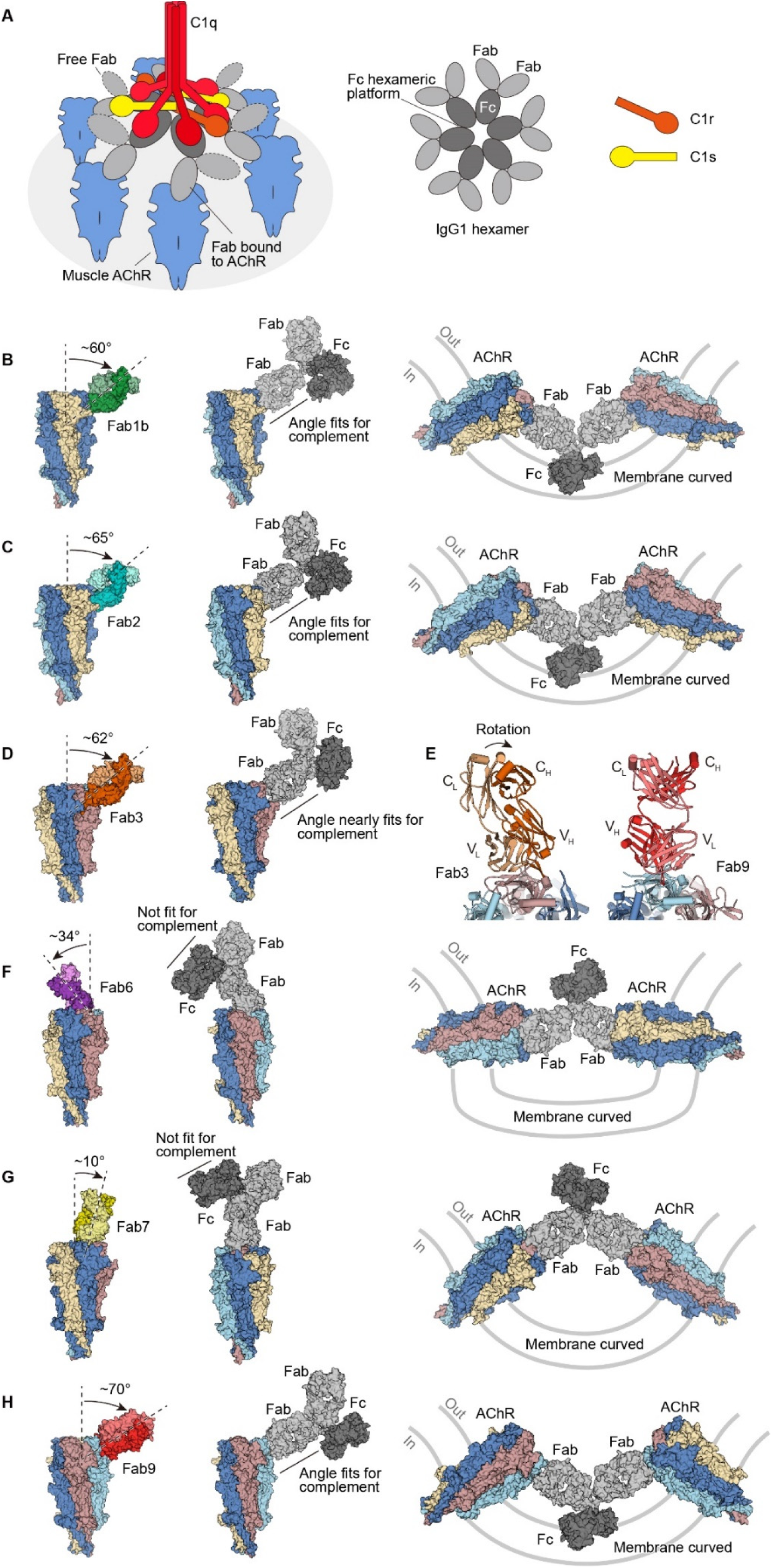
Models for complement activation or receptor internalization related to the angle of Fab binding to the AChR. (A) Cartoon illustration based on recent structures of C1q-mediated complement binding to IgG1 antibodies modeled on AChR Fabs used in this study. (B–D), (F–H) Angles of binding for each Fab were measured against the axis of symmetry down the central pore of the receptor. These Fabs were then used for superposition of an IgG1 structure (PDB 1IGY), which allows for speculation of mechanistic feasibility for either complement activation or curvature of the membrane by cross-linking of receptors. (E) Structure comparison between Fab3 and Fab9 shows that the orientation of Fab3 light and heavy chains has a large rotation when binding to muscle AChR.

**Video S1. Cryo-EM density map quality illustrated by Fab3-bound AChR structure.**

AChR-Fab3 complex shows an example of the high-quality density maps and confidence in defining receptor structure and receptor-autoantibody interactions. The extraordinarily long Fab3 H_V_ CDR3 loop inserts deeply into the ACh site at the α_δ_ and δ interface. Subunit colors are consistent with the main figures.

**Supplemental Data Table 1.**
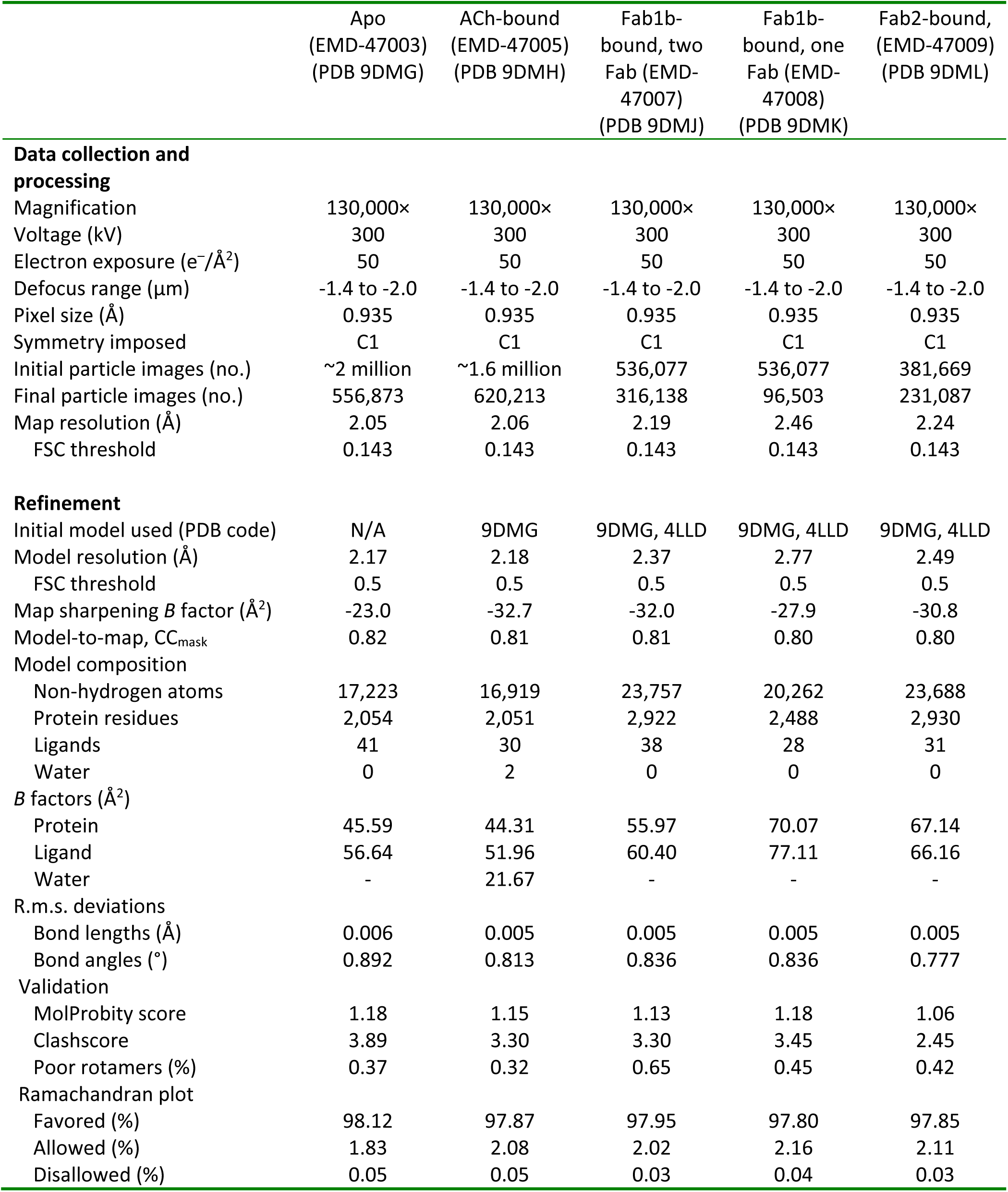
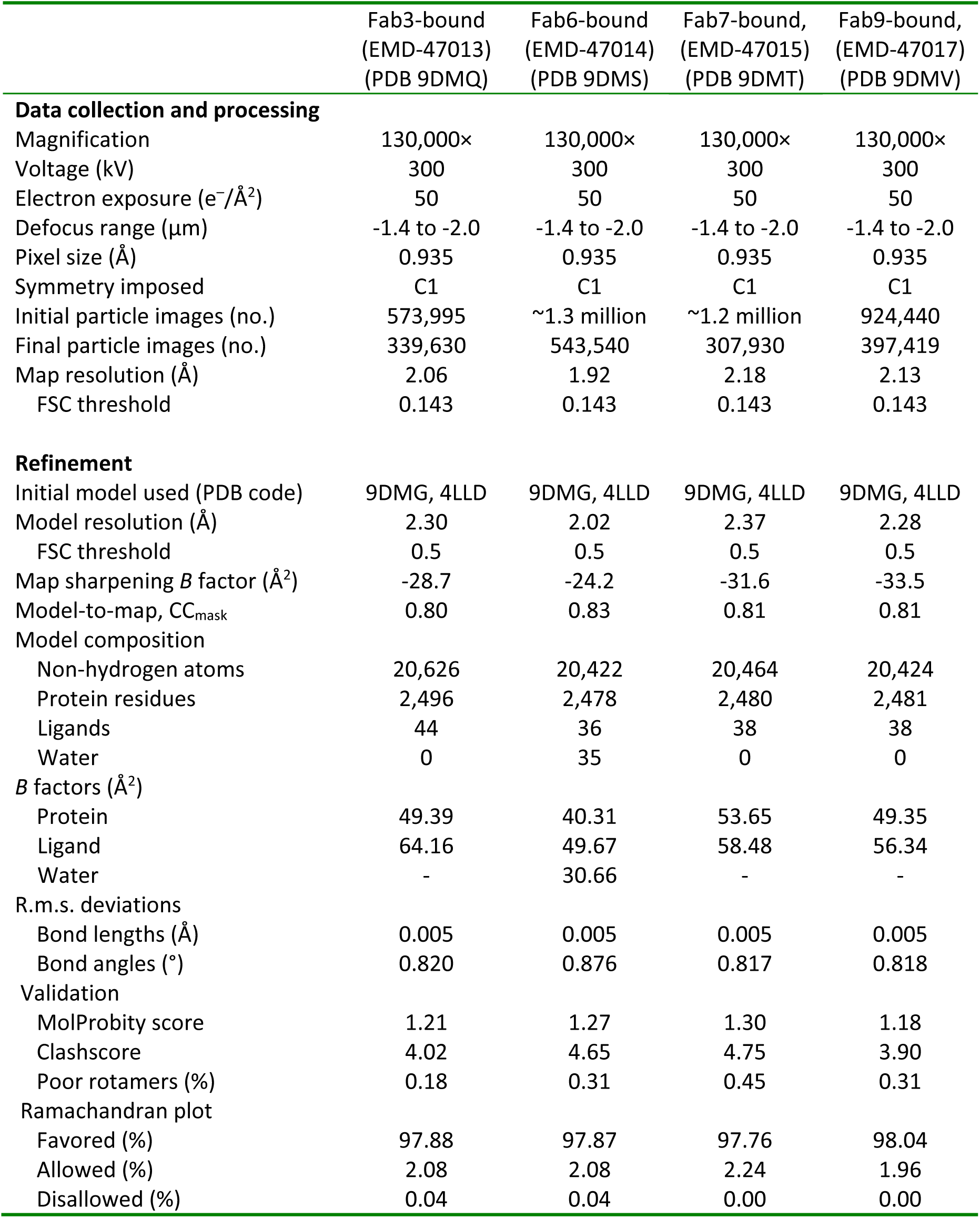
Cryo-EM data collection, refinement and validation statistics.

